# Connexin 43 hemichannels mediate spatial and temporal disease spread in ALS

**DOI:** 10.1101/2020.03.14.990747

**Authors:** Akshata A. Almad, Arens Taga, Jessica Joseph, Connor Welsh, Aneesh Patankar, Sarah K. Gross, Jean-Philippe Richard, Aayush Pokharel, Mauricio Lillo, Raha Dastgheyb, Kevin Eggan, Norman Haughey, Jorge E. Contreras, Nicholas J. Maragakis

## Abstract

Connexin 43 (Cx43) gap junctions and hemichannels mediate astrocyte intercellular communication in the central nervous system under normal conditions and may contribute to astrocyte-mediated neurotoxicity in amyotrophic lateral sclerosis (ALS). Here we show that astrocyte-specific knockout of Cx43 in a mouse model of ALS slows disease progression both spatially and temporally, provides motor neuron (MN) protection, and improves survival. In human ALS tissues and biofluids, we observe that higher levels of Cx43 correlate with accelerated disease progression. Using human iPSC-derived astrocytes (hiPSC-A) from both familial and sporadic ALS, we show that Cx43 is upregulated and that Cx43-hemichannels are enriched at the astrocyte membrane. We then demonstrate that the pharmacological blockade of Cx43-hemichannels in ALS astrocytes, during a specific temporal window, provides neuroprotection of hiPSC-MN and reduces ALS astrocyte-mediated neuronal hyperexcitability. Our data identify Cx43 hemichannels as novel conduits of astrocyte-mediated disease progression and a pharmacological target for disease-modifying ALS therapies.

## Introduction

Astrocytes form a highly coupled intercellular network in the central nervous system (CNS) through gap junctions (GJ) and hemichannels (HC)^1^. GJ facilitate intercellular communication through the exchange of metabolites^2, 3^, ions^4, 5^, second messengers^6, 7^, and even microRNA^8^. Each GJ is composed of two adjoining HC on the plasma membrane of adjacent cells and each hemichannel is made of 6 connexin subunits arranged around a central pore^9^. While connexins mostly form GJ, they can also exit as non-junctional HC that open at the plasma membrane into the extracellular space. Hemichannel opening is dynamic and especially relevant as they open upon cellular damage and under pathological conditions^10–12^. Connexin 43 (Cx43) is the predominant connexin in astrocytes and the major contributor to astrocyte HC^11, 13–15^. Some key roles of Cx43 include^16–19^ homeostatic buffering, synchronization of astrocyte networks, metabolic support for neurons, and modulation of synaptic activity and plasticity, among others.

In amyotrophic lateral sclerosis (ALS), astrocytes carrying disease-specific mutations contribute to disease progression after onset and play a role in motor neuron toxicity^20–22^. ALS astrocyte-mediated neurotoxicity has also been demonstrated *in vitro* and *in vivo* using either murine^22, 23^ or human ALS astrocytes^24–27^. Recent studies suggest specific astrocyte profiles are induced by cytokines that render astrocytes to be proinflammatory and induce toxicity across several neurodegenerative diseases^28^.

Altered Cx43 expression, GJ coupling, and/or HC activity occurs in a number of neurological diseases^10, 11^. In ALS, descriptive studies show Cx43 is increased in the lumbar spinal cord of mutant superoxide dismutase (SOD1^G93A^) mice^29, 30^. In a previous study^31^, we observed an increase in expression of Cx43 in human ALS motor cortex and spinal cord as well as in SOD1^G93A^ mice and that blocking Cx43 HC in SOD1^G93A^ astrocytes is neuroprotective *in vitro*.

This work addresses a central role for astrocytes in ALS and examines their involvement in disease progression. To that end, we utilize an in vivo conditional Cx43KO mouse model to demonstrate that astrocyte Cx43 contributes to the temporal and anatomical progression of disease. We incorporate human brain and spinal cord tissues as well as CSF to show that increases in Cx43 expression are also present in ALS patients and correlate with the rapidity of disease progression. Our fully-humanized, spinal cord specific, platform of SALS and FALS human induced pluripotent stem cell astrocyte (hiPSC-A) and motor neuron (hiPSC-MN) co-cultures allows us to investigate whether blocking Cx43 HC in hiPSC-A using a small molecule has functional relevance in providing neuroprotection and modulating neuronal hyperexcitability to hiPSC-MN as well as other spinal cord neuron subtypes.

## Results

### Astrocyte-specific deletion of Cx43 in SOD1^G93A^ mice slows temporal and anatomical disease progression

As shown previously^31^, Cx43 expression increases in SOD1^G93A^ mice during the course of disease and is accompanied by functional increases in Cx43-mediated GJ and HC activity. To more fully assess the role of astrocyte Cx43 in the context of ALS *in vivo*, we crossed the SOD1^G93A^ mouse with a Cx43fl/fl:GFAP-Cre mouse to specifically knockout Cx43 in astrocytes (**Fig. 1**). Examination of the ventral horn of the spinal cord showed an increase in Cx43 expression in SOD1^G93A^ mice consistent with what we had described previously^31^. However, Cx43fl/fl:SOD1^G93A^:GFAP-Cre (SOD1^G93A^:Cx43 KO) mice show near complete loss of Cx43 expression in the ventral horn as demonstrated by immunofluorescence (**Fig. 1a, b**). This decrease in Cx43 expression is accompanied by a reduction in GFAP expression in the SOD1^G93A^:Cx43 KO mice **(Fig. 1a, b).** In addition, we verified that while Cx43 expression is typically elevated at the symptomatic and endstages, it remains knocked out through endstage (**Fig. 1c)**. The knockout of Cx43 in astrocytes of SOD1^G93A^ mice was anatomically consistent across different segments of spinal cord as demonstrated by Cx43 staining in the cervical and lumbar spinal cord (**Fig. 1c)**.

**Figure 1.**
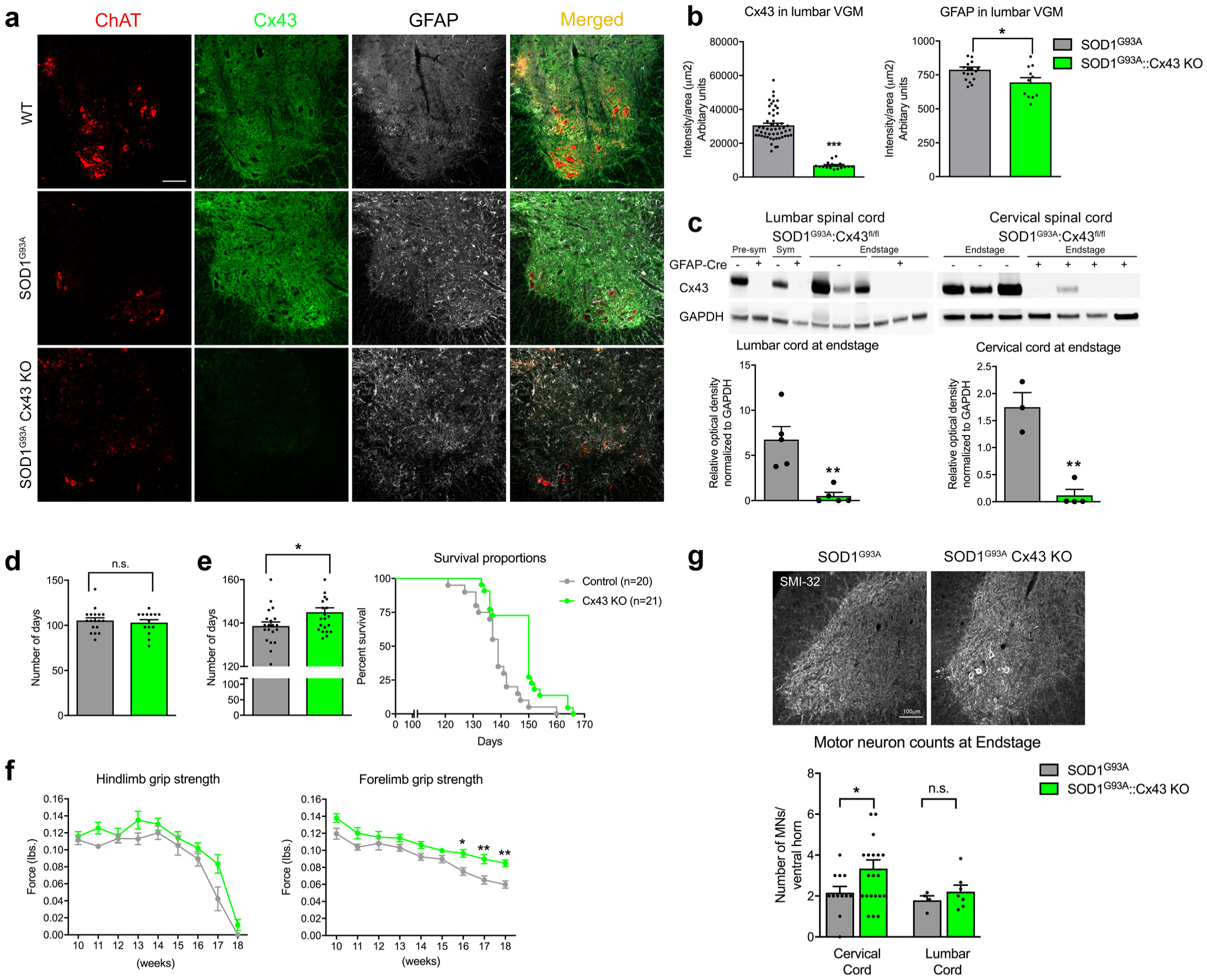
Astrocyte specific deletion of Cx43 improves survival and slows caudal rostral progression in SOD1^G93A^ mice. **(a)** Strategy for conditional deletion of Cx43 in SOD1^G93A^ mice. Cx43^fl/fl^ mice were bred with GFAP-Cre driver line and F1 progeny were further crossed with SOD1^G93A^ mice to generate a triple transgenic mouse. Scale bar=100µm. **(b)** Dramatic reduction in Cx43 occurs in the lumbar spinal cord sections of SOD1^G93A^:Cx43 KO mice compared to SOD1^G93A^ mice. *p<0.05, ***p<0.001, n = 4 animals /group, and at least 3 sections/animal. (**c)** Immunobloting confirmed the deletion of Cx43 in SOD1^G93A^:Cx43^flfl^:: GFAP-Cre+ mice in the lumbar spinal cord at pre-symptomatic, symptomatic and endstage of disease. Cx43 is also absent in cervical spinal cord of the SOD1^G93A^:Cx43^flfl^:: GFAP-Cre+ (n=3-5 animals/group) **p<0.01. **(d)** SOD1^G93A^:Cx43 KO mice along with littermate SOD1^G93A^ mice were followed over time for survival analysis. The onset of the disease was same for both the groups. (**e)** Cx43 KO mice survived significantly longer (145 ± 2.1, n=22) compared to SOD1^G93A^ mice (138.7 ± 1.9, n=20), also depicted in the Kaplan-Meier survival curve, *p<0.05. **(f)** Motor function was tested using grip strength analysis and SOD1^G93A^:Cx43 KO mice displayed significantly higher forelimb grip strength compared to SOD1^G93A^ mice with no overall change in hindlimb grip strength. *p<0.05, **p<0.01. (n=16-20). **(g)** No differences in motor neuron survival was noted at pre-symptomatic and symptomatic stages of disease in the lumbar cord. However, significant preservation of motor neurons was observed at endstage in the cervical spinal cord of SOD1^G93A^:Cx43 KO mice compared to SOD1^G93A^ mice (n=5/group), *p<0.05.

To assess whether the loss of astrocyte Cx43 affects MN survival and disease progression, we performed pathological and motor behavioral studies, specifically grip strength analysis on SOD1^G93A^:Cx43 KO mice during the course of disease. We did not see differences in the timing of hindlimb disease onset in SOD1^G93A^:Cx43 KO mice (**Fig. 1d**), but there was prolongation of survival in the SOD1^G93A^:Cx43 KO mice (**Fig. 1e**). The lack of change in disease onset is also reflected in the motor function using hindlimb grip strength (**Fig. 1f**). While the disease onset started at the same time in all mice, the forelimb grip strength of SOD1^G93A^:Cx43 KO was maintained (**Fig. 1f**). We examined whether this sustained forelimb function and slower disease progression in the SOD1^G93A^:Cx43 KO mice was due to preservation of cervical spinal cord MNs and found a significant preservation of cervical MNs but not lumbar MNs in the SOD1^G93A^:Cx43 KO spinal cord compared to control SOD1^G93A^ spinal cord (**Fig. 1g)**.

Our previous studies^31^ in SOD1^G93A^ mice showed increases in Cx43 but no substantial change in Cx30, the other major astrocyte connexin. However, the SOD1^G93A^:Cx43 KO mice described here did show a modest upregulation of Cx30 in the gray matter, potentially as a compensatory effect from loss of Cx43. We did not see changes in microglial activity by Iba-1 immunostaining and the astrocyte-specific glutamate transporter, GLT-1, was also unchanged, suggesting the Cx43 deletion did not alter the expression of this ALS relevant, astrocyte-specific, glutamate transporter subtype (**Supplementary Fig. 1**).

We also confirmed that the neuroprotection seen *in vivo* was also present using an *in vitro* SOD1^G93A^ astrocyte/WT MN co-culture system^31^. We first validated that SOD1^G93A^ astrocytes exhibit an increase in Cx43 expression, but that the SOD1^G93A^:Cx43 KO lack Cx43 expression as expected (**Supplementary Fig. 2a, b**). Upon culturing these astrocytes with Hb9GFP MN, we demonstrate that SOD1^G93A^ astrocytes induce MN toxicity and this effect is rescued in co-cultures with SOD1^G93A^:Cx43 KO astrocytes resulting in neuroprotection (**Supplementary Fig. 2c**).

**Figure 2:**
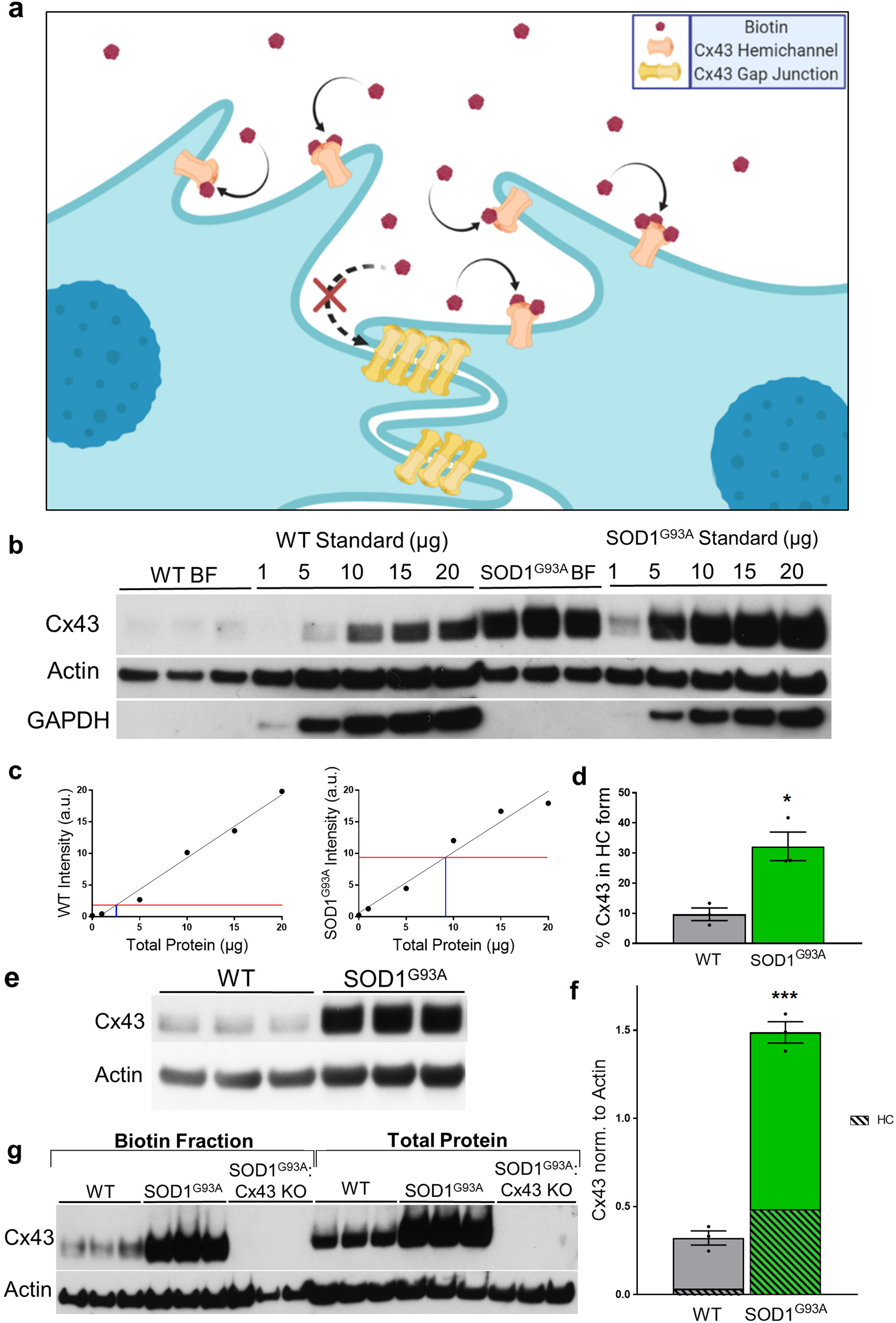
Cx43 HC represents a larger fraction of total Cx43 and is abnormally shifted in the membrane as HC in a mouse model of ALS. **(a)** Graphic representation of biotin binding to Cx43 HC in vitro. In the monolayer system, biotin cannot access and bind to gap junctions, meaning subsequent streptavidin pulldown selects for HC within the biotin fraction. **(b)** Western blot showing Cx43, actin, and GAPDH staining of WT and SOD1^G93A^ biotin fractions (BF) and standards of total protein. Overall Cx43 levels are lower in WT than in SOD1^G93A^. **(c)** The intensities of the WT and SOD1^G93A^ standards were graphed and found to have a linear trend of R^2^ = 0.98 and 0.97. The final calculation predicts WT HC make up 9.6% of total Cx43 *in vitro*, and 32.8% in SOD1^G93A^. **(d)** Individual intensities of all HC were plotted and SOD1^G93A^ HC levels are significantly higher than WT HC levels, (* p < 0.05, t-test, n=3). **(e)** Western blot of WT and SOD1^G93A^ total Cx43 showing significantly higher amounts of total Cx43 in SOD1^G93A^ loaded at 10 µg (quantified in **f**). When mapped as a percentage of total Cx43 identified in b, SOD1^G93A^ Cx43 HC quantification is greater than total Cx43 protein in WT. Significance represents measurement of total Cx43 normalized to actin between the two groups (*** p < 0.001, t-test, n=3). **(g)** Biotin pulldown was repeated with SOD1^G93A^:Cx43 KO samples and total protein to compare against WT and SOD1^G93A^ samples. Negative stains for SOD1^G93A^:Cx43 KO supports full knock-out of Cx43 in all areas or subsets of the cells, and makes the model relevant for HC studies.

Previous work^28, 32^ suggests that activated astrocytes can have either a proinflammatory or “A1” profile that is neurotoxic or an anti-inflammatory “A2” profile that confers neuroprotection. Hence, we determined if potentially SOD1^G93A^:Cx43 KO astrocytes *in vitro* have altered conversion or “activation” compared to WT and littermate SOD1^G93A^ mice based on their transcriptional profiles. However, no consistent pattern was observed to suggest that SOD1^G93A^ astrocytes have an “A1” neurotoxic profile, or that their profile is shifted towards an “A2” neuroprotective phenotype in the SOD1^G93A^:Cx43 KO astrocytes. This suggests that the mechanism of toxicity to MN from SOD1^G93A^ astrocytes and of neuroprotection from SOD1^G93A^:Cx43 KO astrocytes is separate from the shift in astrocyte “activation state” as described by others^28, 32^ (**Supplementary Fig. 3)**.

### Connexin 43 expression is disproportionately shifted to hemichannels in SOD1^G93A^ mouse astrocytes

Having established the role of Cx43 in astrocyte-mediated MN loss and disease progression in murine ALS models, along with our previous observations^31^ that the selective pharmacologic blockade of Cx43 GJ and HC in SOD1^G93A^ astrocytes results in MN protection, we wanted to determine the specific contribution of Cx43 HC to these phenomena. To determine Cx43 HC localization to the astrocyte membrane, we modified a previously described^33^ biotin pulldown assay to specifically target Cx43 HC, which have exposed lysine residues and can bind to biotin in monolayer cultures and therefore allow us to determine what proportion of Cx43 is present as HC (**Fig. 2a**). We first used this pulldown assay on WT and SOD1^G93A^ mouse astrocytes alongside total protein ladders to determine the ratio of HC to total Cx43, and whether there is an absolute increase in Cx43 HC in disease cells (**Fig. 2b**). We then used the Cx43 total protein ladders to create linear regressions for both WT and SOD1^G93A^ cells based on densitometry, and subsequently plotted the individual intensities of the Cx43 stained biotin pulldown lanes against the curve (**Fig. 2c**). Using these curves, we found that HC in WT astrocytes make up 9.6±2.1% of Cx43, whereas HC in SOD1^G93A^ astrocytes are 32.8±4.7% of Cx43 (**Fig. 2d**). While there is an overall increase in Cx43 levels in SOD1^G93A^ (**Fig. 2e**), the absolute levels of Cx43 HC were specifically elevated in disease cells compared to WT astrocytes (**Fig. 2f)**. These data suggest that there is both a significantly larger protein quantity of Cx43 HC present on the surface of SOD1^G93A^ astrocytes than in WT, and that this amount represents an abnormal shift in the percentage of total Cx43 expression to HC formation. We then tested the SOD1^G93A^:Cx43 KO astrocytes against WT and SOD1^G93A^ astrocytes as a negative control and found that the Cx43 KO cells do not have a measureable quantity of either total Cx43 or Cx43 in HC form (**Fig. 2g**).

### Connexin 43 expression in sporadic ALS tissues and biofluids correlates with the severity of disease progression

In our published work^31^, we demonstrated an increase in Cx43 expression in ALS patient motor cortex and spinal cord. Here, we have expanded the cohort of SALS motor cortex and cervical spinal cord **(Table 1)** and examined the correlation between Cx43 expression and the temporal course of disease. First, we observed an increase in Cx43 and GFAP transcript levels in the motor cortex of post mortem SALS patients (**Fig. 3a, b).** This increase in Cx43 transcript parallels protein expression levels, where Cx43 expression is increased in both ALS motor cortex and cervical spinal cord when compared to controls. Furthermore, patients with rapidly progressing disease (i.e. deceased within 2 years after onset)^34, 35^ display higher Cx43 protein levels in both motor cortex and cervical spinal cord compared to patients with a more typical disease progression (i.e. deceased between 2 −5 years after onset)^34, 35^ (**Fig. 3c, d**). The correlation between a faster temporal course of disease and Cx43 expression in ALS tissue was also paralleled by CSF samples from ALS patients. We observed a trend towards an inverse correlation between CSF Cx43 levels and the ALS Functional Rating Scale-Revised (ALSFRS-R)^34, 35^ (**Fig. 3e**). The Cx43 protein level was robustly elevated in the CSF samples obtained from ALS patients with rapid progression, compared to slower progressing patients (**Fig. 3f**).

**Figure 3:**
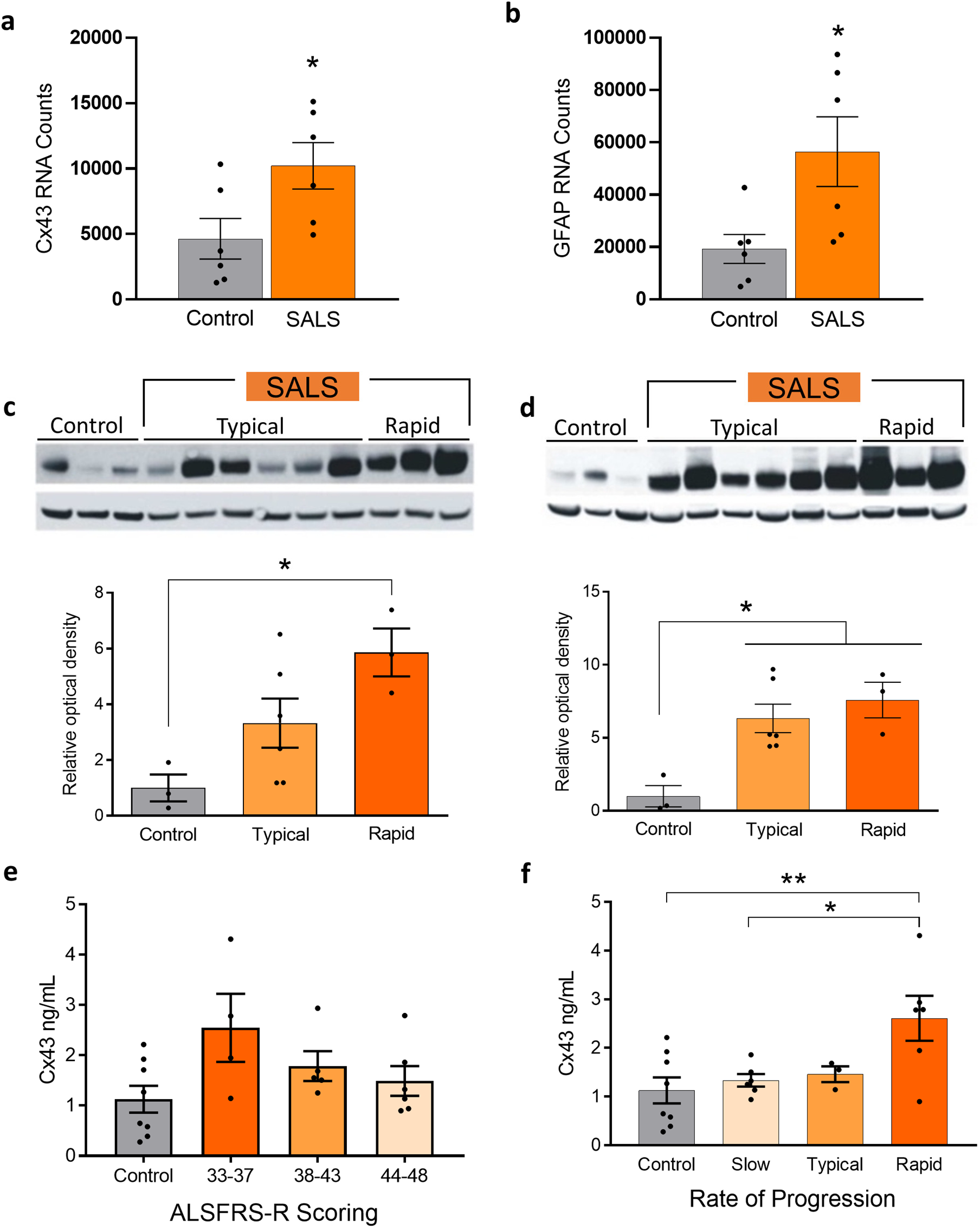
Cx43 transcription and expression in human post-mortem tissue and CSF correlates with rapidity of disease progression in ALS. **(a)** NanoString analysis of Cx43 motor cortex from SALS patients and controls shows significant increase (* p < 0.05, t-test, n=6/condition). **(b)** Similarly, NanoString analysis of GFAP in motor cortex shows significant changes (* p < 0.05, t-test, n=6/condition). **(c)** Post-mortem motor cortex tissue shows significant increase of Cx43 by Western blot of fast sporadic patients compared to healthy controls. Quantification pictured below (* p < 0.05, one-way ANOVA, n=3-6/condition). **(d)** In both typical and fast sporadic patient cervical cords, Cx43 is significantly higher. Quantification pictured below (* p < 0.05, one-way ANOVA, n=3-6/condition). **(e)** CSF from control and sporadic patients based on ALSFRS-R scores shows a trend towards increased Cx43 in CSF with lower scores (n=4-8/condition. **(f)** CSF from patients and healthy volunteers shows significantly higher levels of Cx43 by ELISA in fast sporadic patients compared to control and slow progression patients all other progression rates (** p < 0.01, * p < 0.05, one-way ANOVA, n=3-8/condition). All error bars represent SEM.

**Table 1.** Control and SALS autopsy tissue samples used in this study.

| VA Tissue Bank # | Age of death | Gender | Category | Cause of Death |
| --- | --- | --- | --- | --- |
| 100012 | 81 | Female | Control | Respiratory insufficiency; acute colitis |
| 110005 | 62 | Male | Control | Acute pulmonary edema |

| 110006 | 68 | Male | Control | Bronchopneumonia;<br>lymphoproliferative disorder |
| --- | --- | --- | --- | --- |
| Target<br>ALS # | Age of<br>death | Approximate disease<br>duration (months) | Diagnosis | C9ORF72 |
| 93 | 68 | 30 | SALS | No |
| 98 | 78 | 28 | SALS | N/A |
| 102 | 72 | 28 | SALS | N/A |
| 104 | 71 | 25 | SALS | No |
| 13 | 71 | 132 | SALS | No |
| 3 | 44 | 192 | SALS | No |
| 49 | 68 | 70 | SALS | No |
| 97 | 68 | 71 | SALS | No |
| 3 | 77 | 14 | SALS | No |
| 32 | 59 | 18 | SALS | No |
| 89 | 71 | 22 | SALS | No |
| 9 | 59 | 24 | SALS | No |

### Astrocyte Cx43 HC expression and function is increased in both FALS and SALS patients

We differentiated human induced pluripotent stem cell-derived astrocytes (hiPSC-A) using a spinal cord patterning protocol^36^ (**Supplementary Fig. 4**) and examined Cx43 expression in a large cohort of patients with SALS and FALS (**Table 2**). Since the differentiation of hiPSC-A occurred in the absence of neurons, we anticipated that any Cx43 changes would be independent of neuronal input (cell autonomous) and not as a secondary effect of neuron-dependent astrocyte “activation”. A consistent and significant increase in Cx43 expression was observed in hiPSC-A from SALS patients (**Fig. 4a)**. In addition, we examined FALS hiPSC-A, specifically from 3 different SOD1^D90A^ and 3 unrelated SOD1^A4V^ patients. While the expression of Cx43 in the SOD1^D90A^ samples had a trend towards an increase in Cx43 expression, a significant increase in Cx43 expression occurred in the SOD1^A4V^ hiPSC-A compared to control hiPSC-A (**Fig. 4b**). Interestingly, Cx43 levels in hiPSC-A had a direct correlation with ALS disease progression: Cx43 was higher in patients with SOD1^A4V^ mutation, who exhibit a faster disease progression, while Cx43 expression increased but not significantly in SOD1^D90A^ cases, who have a relatively slower temporal course^37^. To examine if correction of the SOD1^A4V^ mutation attenuated Cx43 expression, we used a previously^38^ characterized ALS SOD1^A4V^ hiPSC line and its gene corrected isogenic control line (**Supplementary Fig. 5a**). As expected, Cx43 protein levels were higher in the ALS SOD1^A4V^ hiPSC-A compared to control astrocytes and the corresponding gene corrected SOD1^A4V^ line showed a partial normalization of Cx43 expression (**Supplementary Fig. 5b**).

**Figure 4:**
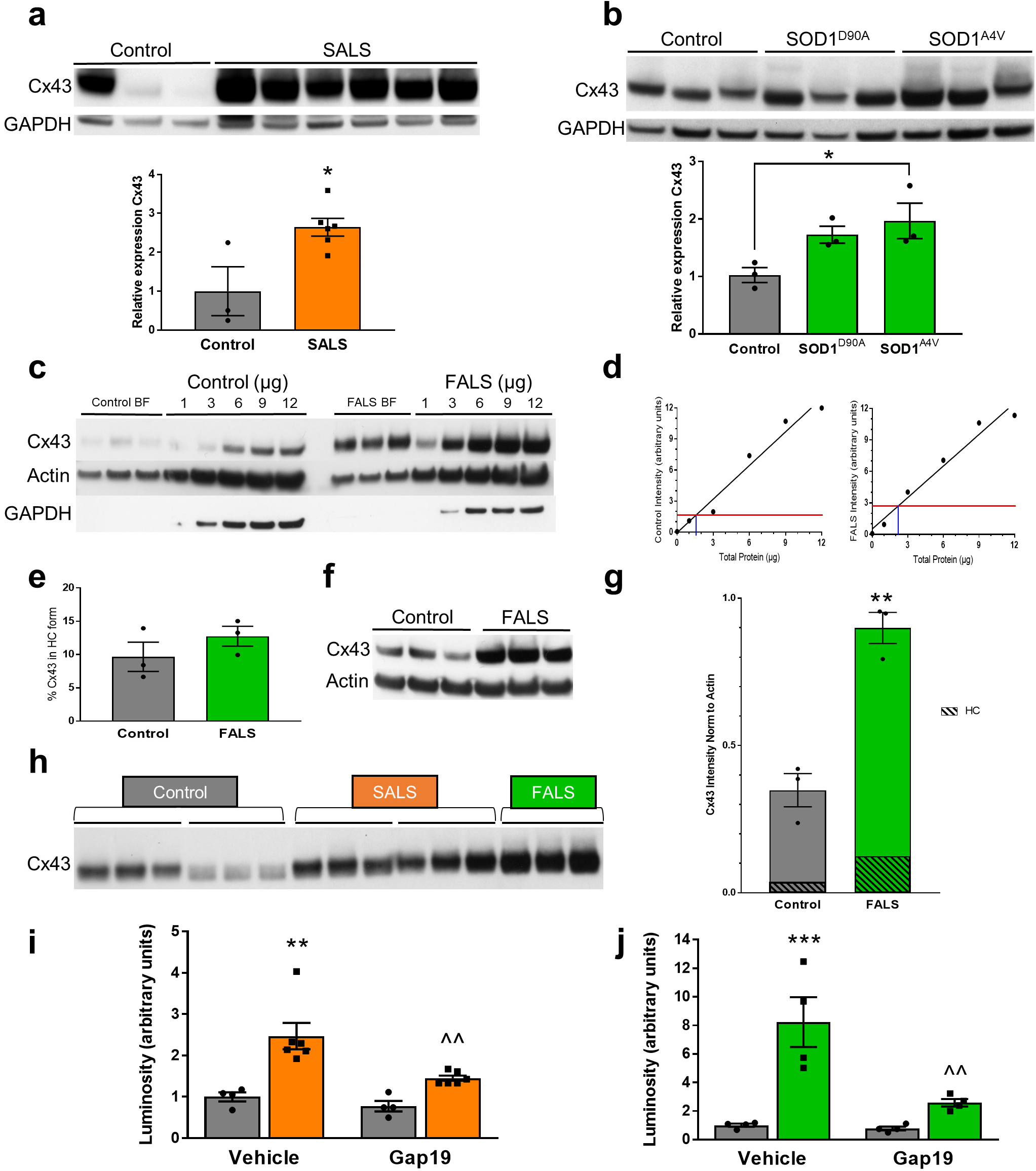
Human ALS iPSC-Astrocytes have increased Cx43 HC quantities associated with increased cell permeability which can be blocked by Gap19. **(a)** Western blots of total protein collected from various human iPSC-A control and sporadic ALS lines were compared for Cx43 expression. Quantification shows sporadic Cx43 expression is significantly higher (* p < 0.05, t-test, n=3-6/condition). **(b)** Similarly, Western blots stained for Cx43 in hiPSC-A from control and familial lines shows that in A4V patients, Cx43 expression is significantly higher (* p < 0.05, one-way ANOVA, n=3/condition). **(c)** Using the same protocol as described in Fig. 2, control and familial hiPSC-A in vitro lines were subjected to biotin pulldown and subsequent biotin fractions and total protein were blotted for Cx43 by Western blot, with actin and GAPDH as housekeeping genes. **(d)** Linear quantification against the total protein standards have R^2^ values of 0.97 and 0.96. **(e)** There is not a statistically significant shift in the percent of Cx43 as HC between ALS and control hiPSC-A. **(f)** Western blot of total protein for hiPSC-A control and FALS lines. **(g)** FALS has significantly more Cx43 than control (** p < 0.01, n=3). When percentages from 4e are mapped against total protein, FALS has more Cx43 in HC format by absolute calculation than control. **(h)** Western blot of two lines from control and SALS, and one line of FALS patients stained for Cx43 from the biotin fraction *in vitro*. Each bar represents one line tested in triplicate. **(i)** ATP levels detected in the supernatants of control and sporadic hiPSC-A show significantly higher amounts in sporadic astrocytes compared to controls. Additionally, treatment with Gap19, a HC specific blocker, significantly reduces supernatant ATP in sporadic astrocytes (two-way ANOVA, ** p < 0.01, ^^ p < 0.01, n = 4-6). **(j)** Similarly, supernatant ATP of familial hiPSC-A is significantly higher than control without treatment. Gap19 also significantly reduces supernatant ATP of familial astrocytes compared to vehicle treatment (two-way ANOVA, *** p < 0.001, ^^ p < 0.01, n = 4).

**Table 2.** Human iPSC-derived astrocyte and MN lines used in this study.

| hiPSC astrocyte line | Phenotype |
| --- | --- |
| CS25 | Control |
| CiPS | Control |
| JH082 | Control |
| GO018 | Control |
| JH013 | Slow Sporadic |
| JH036 | Slow Sporadic |
| JH029 | Typical Sporadic |
| JH017 | Typical Sporadic |
| JH058 | Rapid Sporadic |
| JH040 | Rapid Sporadic |
| GO017 | Slow Familial (SOD1<sup>D90A</sup>) |
| GO004 | Slow Familial (SOD1<sup>D90A</sup>) |
| GO 028 | Slow Familial (SOD1<sup>D90A</sup>) |
| GO013 | Rapid Familial (SOD1<sup>A4V</sup>) |
| GO 002 | Rapid Familial (SOD1<sup>A4V</sup>) |
| GO 007 | Rapid Familial (SOD1<sup>A4V</sup>) |
| 39B | Rapid Familial (SOD1<sup>A4V</sup>) |
| 39B2.5 | 39B isogenic corrected line (SOD1<sup>+/+</sup>) |
| hiPSC-MN line | Phenotype |
| GM01582 | Control |
| CS25 | Control |

Similar to murine *in vitro* studies, we determined the specific contribution of Cx43 HC to the amount of total Cx43 using ALS hiPSC-A. We used the biotinylation assay in FALS hiPSC-A (**Fig. 4c-g**) as described in **Fig. 2**. Based on linear regression analysis, both control and the FALS hiPSC-A expressed similar proportional amounts of Cx43 as HC (**Fig. 4d**). Therefore, when quantified, the percentages of Cx43 present as HC were not significantly different (**Fig. 4e**). However, when we blotted for total Cx43 protein (**Fig. 4f**), FALS had higher absolute levels of Cx43 expression (**Fig. 4g**). We extended our analysis to the absolute amounts of Cx43 HC between pulldowns from multiple control and SALS patient lines, and noted that all ALS hiPSC-A lines had higher amounts of Cx43 HC present on their membranes (**Fig. 4h**). We next tested the opening of Cx43 HC by detecting ATP via a luciferase assay in the supernatants of FALS and SALS hiPSC-A. Endogenous ATP in this instance acts as a functional marker for HC opening. At baseline, both FALS and SALS hiPSC-A had significantly higher concentrations of ATP in the supernatant. Upon treatment with Gap19, a Cx43 HC specific mimetic peptide blocker^39^, control ATP supernatant levels were not reduced but ATP, in both the FALS and SALS hiPSC-A supernatants, was reduced to within control ranges (**Fig. 4i, j**). This suggests that, at baseline, ALS hiPSC-A have increased HC permeability to ATP that can be pharmacologically blocked.

### Human iPSC-A Cx43 HC mediate iPSC-MN toxicity in both sporadic and familial ALS

We have previously shown that the addition of Cx43 GJ and HC-specific blockers resulted in neuroprotection in mouse SOD1^G93A^ astrocyte and WT MN co-cultures^31^. Given the opportunity to examine the genetic and phenotypic heterogeneity of ALS using hiPSC-A, we co-cultured hiPSC-A (**Fig. 5a**) from control and 2 unrelated SOD1^A4V^ ALS patients with hiPSC-MN from controls. We found that SOD1^A4V^ hiPSC-A induced toxicity in human iPSC-MN (ChAT^+^), including a subset of Isl1^+^ motor neurons, and that blocking Cx43 HC with Gap19 provided significant neuroprotection (**Fig. 5a-k**). Similarly, we employed an identical co-culture paradigm to examine whether hiPSC-A from SALS patients would be toxic to hiPSC-MN (**Fig. 5a**). We observed a similar level of hiPSC-MN toxicity compared to the SOD1^A4V^ hiPSC-A and also demonstrated neuroprotection via hemichannel blockade using Gap19 (**Fig. 5a-k**). Interestingly, when the SOD1^A4V^ mutation was corrected using zinc finger endonuclease (**Supplementary Fig. 5**), the resulting isogenic line displayed residual neurotoxicity on hiPSC-MN. This neurotoxicity could be further rescued by Cx43 HC blockade with Gap19 (**Supplementary Fig. 5c-m**).

**Figure 5:**
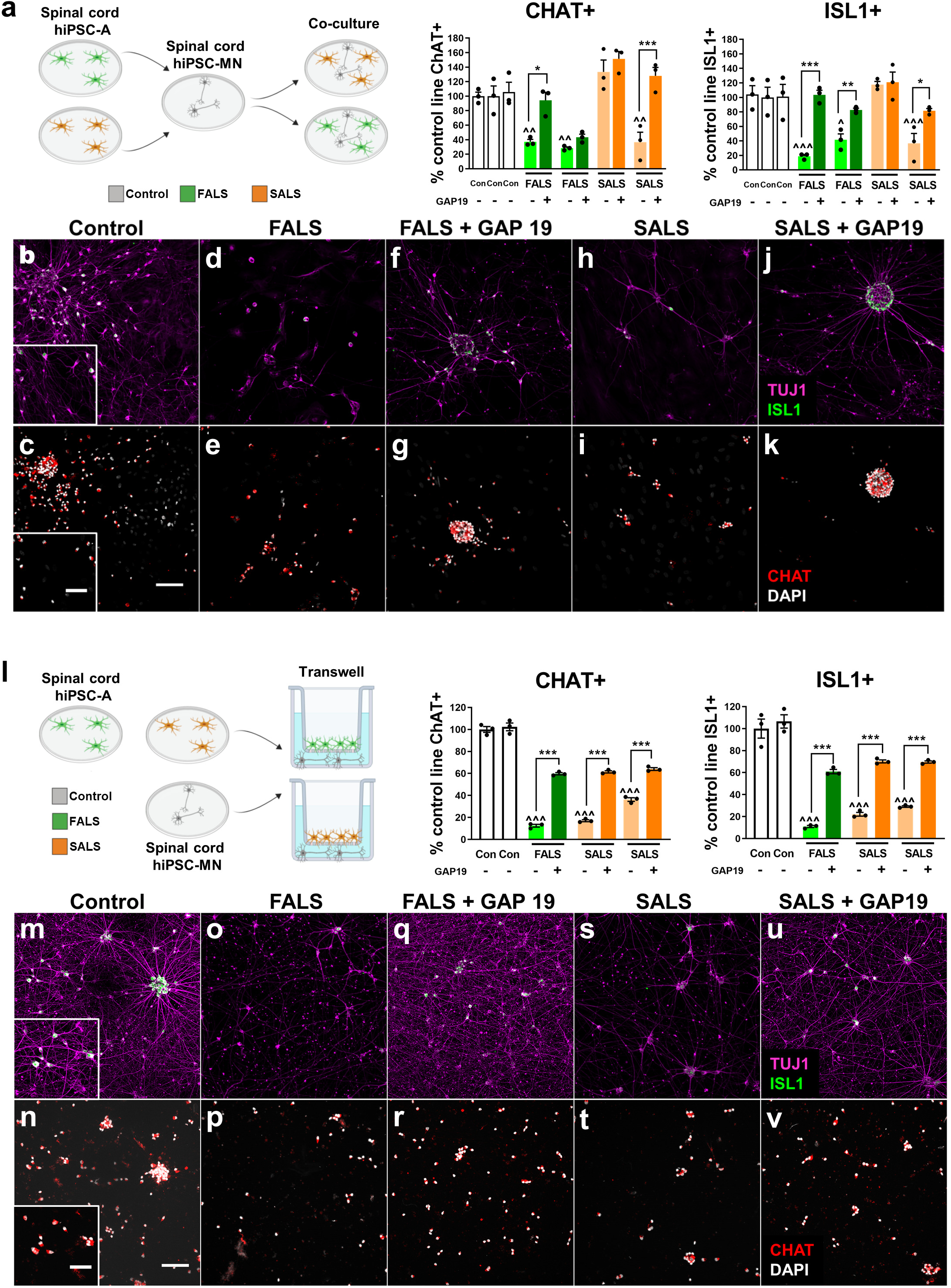
Human iPSC-MN toxicity is mediated by Connexin 43 HC from familial as well as sporadic ALS hiPSC-A. **(a-k)** Motor neurons derived from control hiPSC were plated with hiPSC-A from either control, SALS or FALS (SOD1^A4V^) patients. The number of hiPSC-MN (ChAT^+^ and Isl-1^+^) plated on familial and sporadic ALS hiPSC-A decreased significantly over a period of 14 days compared to motor neurons plated with control hiPSC-However, when co-cultures containing ALS hiPSC-A were treated with a Cx43 hemichannel blocker (Gap19), a significant increase in hiPSC-MN survival was observed. **(l-v)** The co-culture of hiPSC-A/-MN in a transwell system allowed neurons and astrocytes to share the same medium, while preventing direct cell contact. FALS (SOD1^A4V^) and SALS hiPSC-A shows ChAT^+^ and Isl-1^+^ hiPSC-MN toxicity rescued by Gap19. Bars represent the means±SEM with data points of n=3 coverslips per condition. Significant comparisons (one-way ANOVA) between untreated control and ALS co-cultures are marked with (^), while significant effects of GAP19 on co-cultures containing ALS astrocytes are marked with (*). * or ^ p<0.05; ** or ^^ p<0.01; *** or ^^^ p<0.001. Scale bar=50µm, inset scale bar=20 µm.

In order to confirm our findings of Cx43 HC-specific neurotoxicity, we first established that ALS hiPSC-A toxicity was conferred by a factor released into the medium, as might be expected for connexin HC. To this end, we performed a hiPSC-A/MN co-culture study where hiPSC-A were grown on a transwell insert above and control hiPSC-MN on the base of the well, with both cell groups grown in the same media (**Fig. 5l**). We co-cultured FALS and SALS hiPSC-A lines with hiPSC-MN derived from control patients and observed a significant reduction in hiPSC-MN survival as defined by ChAT and Isl1+ immunostaining, similar to the direct co-culture paradigm (**Fig. 5l-v**). We also treated the transwell co-cultures with Gap19, which resulted in significant neuroprotection from both FALS hiPSC-A as well as SALS hiPSC-A induced toxicity (**Fig. 5l-v**). This suggests that astrocyte-mediated toxicity from these ALS lines is propagated mainly through HC. Similar to the co-culture paradigm, the correction of the SOD1^A4V^ mutation in the FALS hiPSC-A line (**Supplementary Fig. 5**) conferred partial neuroprotection, with the residual neurotoxicity being further rescued by Cx43 HC blockade with Gap19 (**Supplementary Fig. 5n-x)**. We confirmed that the neuroprotective effect of Gap19 was specifically mediated by astrocytes, as the addition of Gap19 to hiPSC-MN cultures alone did not affect MN survival (**Supplementary Fig. 6**).

While much of ALS *in vitro* biology is focused on MN loss, there is *in vivo* evidence^40–42^ that other neuronal populations are affected by neurodegeneration. We utilized our hiPSC-based platform to examine whether non-motor neuron degeneration in ALS is astrocyte and specifically Cx43 HC mediated. Our data utilizing human ALS hiPSC-A/MN co-cultures show that ALS astrocytes from both SALS and FALS patients induce toxicity to ChAT^-^/TUJ1^+^ neurons. This effect was partially rescued by the addition of the Cx43 HC blocker Gap19 (**Supplementary Fig. 7a**). We established, by using the transwell system and Gap19 as described above, that this ALS astrocyte-mediated neurotoxicity was related to a factor released through Cx43 HC (**Supplementary Fig. 7b**). Our findings on astrocyte and Cx43 HC mediated hiPSC-A neurotoxicity towards MN and non-MN populations could also be reproduced using a different control hiPSC-neuronal line (CS25) (**Supplementary Fig. 8**).

### The Cx43 HC blocker tonabersat provides dose-dependent neuroprotection to hiPSC-MN

Our findings from rodent models and from human ALS samples suggest that Cx43 HC may provide an appropriate target for ALS therapeutics. Tonabersat (SB-220453) was selected for its translational potential in human clinical trials and has been shown^43, 44^ to specifically block Cx43 HC at low concentrations (1-10 μM).

To examine the potential of tonabersat in preventing hiPSC-A and Cx43 HC mediated neurotoxicity, we co-cultured control hiPSC-MN with either control hiPSC-A, FALS hiPSC-A, or SALS hiPSC-A. We utilized direct co-culture (**Fig. 6a-k**) and transwell (**Fig. 6l-v**) paradigms as described above and found that tonabersat protects hiPSC-MN from ALS astrocyte-mediated death in a dose-dependent manner in both systems. At doses of 1µM and 10µM, tonabersat blocks Cx43 HC and rescues ChAT^+^ MNs, including a subpopulation of Isl1^+^ MN. A higher concentration of 100µM, tonabersat was not neuroprotective, and likely reflects a long-term effect on reducing gap junction trafficking instead of HC activity, as described previously^43^ (**Fig. 6a-k**). Interestingly, we also observed that tonabersat was protective to non-MN populations (ChAT^-^ and Isl1^-^ neurons), similar to the results seen with Gap19 treatment (**Supplementary Fig. 9a**). The neuroprotective effect of tonabersat was also seen in a transwell study (**Fig. 6l-v**), suggesting that tonabersat’s neuroprotection is HC-mediated and further confirming the importance of Cx43 HC in toxicity. This HC-mediated neuroprotection was also observed with non-MN subtypes (**Supplementary Fig. 9b**). We believe that the neuroprotective effects of tonabersat are specific to Cx43 HC on astrocytes since the application of tonabersat to cultures of hiPSC-MN alone did not have any effect on hiPSC-MN survival (**Supplementary Fig. 10**).

**Figure 6:**
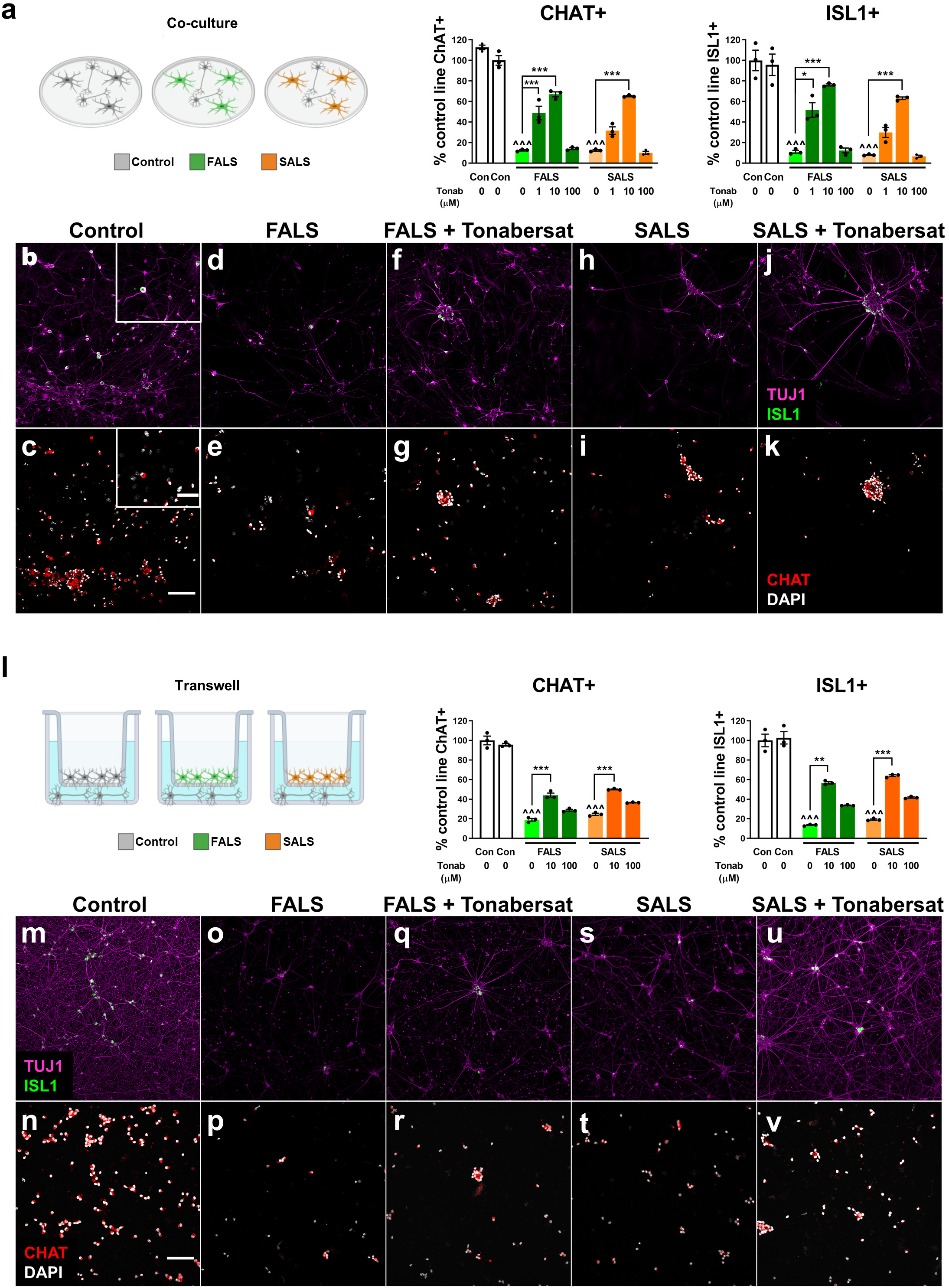
The Cx43 HC blocker tonabersat provides dose-dependent neuroprotection to hiPSC-MN. **(a-k)** FALS and SALS hiPSC-A/MN co-culture immunostained for ChAT and Isl-1 shows dose-dependent neuroprotection from tonabersat (1µM, 10µM and 100µM), a small molecule that acts as Cx43 blocker, after a 14 day incubation period. **(l-v)** Following transwell co-culture of FALS and SALS hiPSC-A with hiPSC-MN, immunostainining for ChAT^+^ MN and Isl1^+^ MN confirms dose-dependent neuroprotection with tonabersat. Significant comparisons (one-way ANOVA) between untreated control and ALS co-cultures are marked with (^), while significant effects of tonabersat on co-cultures containing ALS astrocytes are marked with (*). * or ^ p<0.05; ** or ^^ p<0.01; *** or ^^^ p<0.001, n=3/condition Scale bar=50µm, inset scale bar 20µm.

In the CNS, tonabersat’s actions may translate to an astrocyte-mediated reduction of neuronal excitability^45, 46^, as suggested by *in vivo* studies showing tonabersat’s inhibitory effects on cortical spreading depression^47, 48^ and seizures^49, 50^. We have previously used^36^ multielectrode array (MEA) in a fully human iPSC-based platform to evaluate how astrocytes influence motor neuron electrophysiology (**Supplementary Fig. 11a**), and here we employ that system to evaluate the electrophysiological actions of tonabersat *in vitro*. We first utilized control co-cultures of hiPSC-A/MN on MEA plates to study how the pharmacological blockade of hiPSC-A Cx43 HC would affect hiPSC-MN electrophysiological activity. The addition of tonabersat (**Supplementary Fig. 11b**) at a concentration of 10 µM significantly reduced neuronal spiking and bursting activity within 5 minutes of application. The effect was dose-dependent, as a lower concentration of tonabersat (i.e. 1 µM) was less effective (**Supplementary Fig. 11b**). These actions parallel those of Cx43 HC-specific blocker Gap19 (**Supplementary Fig. 11c**), confirming our previous observations that Cx43 HC influences neuronal firing^36^. To ensure that these effects were astrocyte-mediated, we tested tonabersat on hiPSC-MN alone in culture and did not observe any change in neuronal electrophysiological activity (**Supplementary Fig. 11d**).

We then evaluated these electrophysiological findings in the context of ALS (**Fig. 7**). Human control and ALS hiPSC-A were co-cultured with control hiPSC-MN and then recorded serially for 2 weeks. We observed a transient increase in neuronal spiking and bursting activity in the co-cultures with SALS and FALS hiPSC-A when compared to control co-cultures (**Fig. 7a**), which occurred early during the 2-week recording period (most significantly at DIV 6 of co-culture), followed by a depression of electrophysiological activity (particularly at DIV 14 of co-culture). This transient ALS astrocyte-mediated hyperexcitability was significantly reduced by the acute treatment with tonabersat (**Figure 7b**), to levels of spiking and bursting activity not significantly different from those of the untreated control co-cultures.

**Figure 7.**
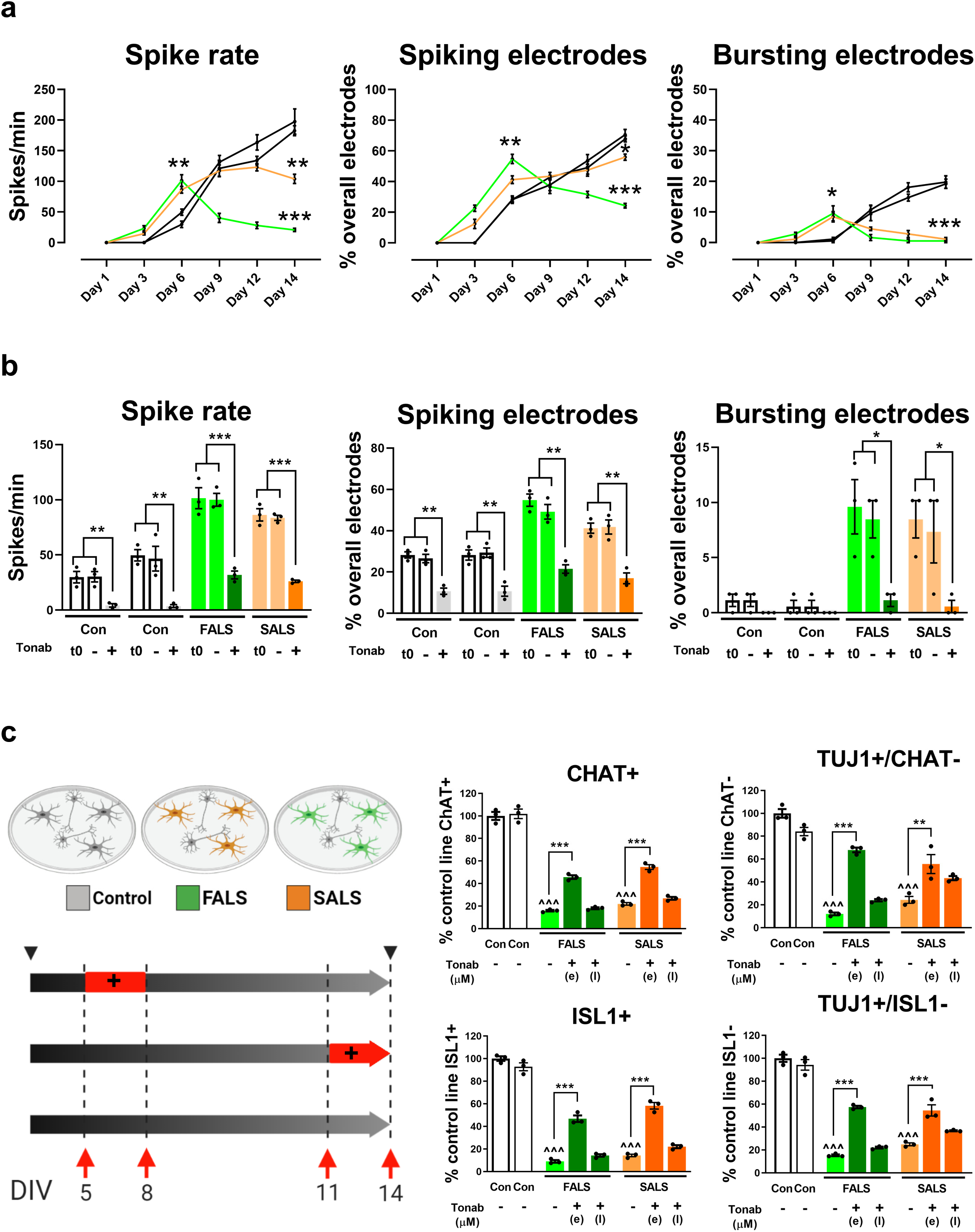
Multielectrode array recordings (MEA) from hiPSC-Astro/MN cultures demonstrate that tonabersat Cx43 HC-mediated effects on hiPSC-MN electrophysiology correlates with its neuroprotective actions. **(a)** Multieledrode array recording of neuronal activity (spike rate, percentage of overall MEA electrodes displaying spiking or bursting activity) over-time from co-cultures between control hiPSC-MN and control or ALS hiPSC-A. The presence of SALS and FALS astrocytes when compared to control astrocytes determines early increases in spiking and bursting activity followed, at later time points, by reduced electrophysiological activity. Most significant time-point comparisons (i.e. day 6 and day 14 of co-culture) between control and ALS conditions are marked with (*). Data are mean±SEM (mean of n = 3 MEA plates per condition, one-way ANOVA). **(b)** MEA activity within 5 minutes after the application of 10µM tonabersat (+) on co-cultures with control or ALS hiPSC-A compared to baseline (t0) and vehicle (-). MEA baseline activity is day 6 time point shown in Figure 7a. Tonabersat results in significant inhibition (*) of neuronal spiking and bursting activity. **(c)** The effects of 10µM tonabersat (+) on hiPSC-neurons survival in control and ALS co-cultures was tested as outlined in figure 6, but for a shorter time course of 3 days, either early (DIV 5 to 8, or “e”) or late (DIV 11-14 or “l) during co-culture. Significant neuroprotection was appreciated after early but not late exposure to tonabersat, and was evident for both motor neuronal (ChAT+ and ISL1+ cells) and non-motor neuronal cell types (TUJ1+/ChAT-, TUJ1+/ISL1+). Significant comparisons (one-way ANOVA) between untreated control and ALS co-cultures are marked with (^), while significant effects of tonabersat on co-cultures containing ALS astrocytes are marked with (*). * or ^ p<0.05; ** or ^^ p<0.01; *** or ^^^ p<0.001, n=3/condition

We then verified whether this ALS astrocyte-mediated neuronal hyperexcitability correlated temporally with our previous findings on ALS astrocyte and Cx43 HC mediated neurotoxicity. To address this, we used the co-culture paradigm with ALS and control hiPSC-A/MN co-cultures but with an abbreviated course of treatment with tonabersat (**Figure 7c**). Interestingly, we observed neuroprotection of control hiPSC-MN in co-culture with ALS hiPSC-A only when 10 µM tonabersat was applied at an early-time point (i.e. from day 5 to day 8 of co-culture), which corresponded to the period of neuronal hyperexcitability as recorded by MEA. This protection did not occur at a later time-point (from day 11 to day 14 of co-culture), when MEA demonstrated neuronal hypoactivity in co-cultures with ALS hiPSC-A. These effects were not limited to motor neurons, as we observed similar neuroprotection for ChAT^-^ neurons as well (**Fig. 7c**).

## Discussion

Connexin 43 is the predominant connexin in astrocytes^11, 13–15, 51^ and is unique as it is the only astrocyte connexin that can be incorporated into hemichannels as well as gap junctions. Changes in Cx43 expression are reported in neurodegenerative disorders, primarily as an upregulation of the protein^31, 52–55^, although less is known about the degree to which these increases are incorporated specifically into HC, GJ or both^56^.

In ALS, a few studies have shown increased total Cx43 in lumbar spinal cord of SOD1^G93A^ mice, although the implications for the increases in total Cx43 protein were not examined^29, 30, 57^. In our previous study^31^, we observed an increase in expression of Cx43 in this ALS mouse model. While the upregulation of Cx43 could be attributed to an astrocytic response to neuronal cell death *in vivo*, our recent work suggests that the increased expression of Cx43 in SOD1^G93A^ mice has a cell autonomous component, since SOD1^G93A^ astrocytes cultured *in vitro* in the absence of neurons or neuronal death displayed increases in Cx43 expression^31^. These increases in astrocyte Cx43 has pathophysiological relevance since the pharmacological blockade of Cx43 in SOD1^G93A^ astrocytes can partially protect MN from astrocyte-mediated neurotoxicity.

Astrocytes have been implicated in ALS progression after disease onset^21^. In order to investigate astrocyte Cx43-specific contributions to disease progression, we generated a novel triple transgenic SOD1^G93A^ mouse with Cx43 KO in astrocytes using GFAP-Cre. While the disease onset was not affected by astrocyte-specific Cx43 KO, we observed a delayed decline of forelimb motor function and an overall increase in survival in this mouse model. These observations suggest that astrocyte Cx43 influences both the temporal disease course after onset and the anatomical progression of disease. The alteration of the temporal and anatomical progression following the knockout of astrocyte Cx43 and Cx30 has similarly been reported in a spinal cord injury model^58^. Previous studies in Alzheimer’s disease, however, have suggested that GJ coupling may be a mechanism for compensatory neuroprotection by buffering potentially toxic substances through the astrocyte network and away from at-risk neurons^59, 60^. Therefore, it may be that the KO of Cx43, which we show affects both GJ and HC formation in our SOD1^G93A^:Cx43 KO mice, may actually underestimate the potential neuroprotective effect of Cx43 HC loss alone. This would be consistent with our previous data in SOD1^G93A^ *in vitro* studies showing that pharmacologic blockade of Cx43 HC provided equivalent neuroprotection to the less specific pharmacological blockade of both Cx43 GJ and HC^31^. The modest upregulation of Cx30, as a GJ protein, in the ventral gray matter could possibly contribute to neuroprotection as well. Cx43 HC opening has also been implicated in altering astrocyte morphology and triggering cell death in both astrocytes and neurons^61, 62^.

Previous work^28^ has suggested that activated microglia release specific factors that result in the generation of toxic A1 astrocytes while a second population of A2 astrocytes are neuroprotective. Early data have suggested that these A1 phenotypes may have a role in neurodegenerative diseases^28, 32^. We found that SOD1^G93A^ mouse astrocytes responsible for MN toxicity did not exhibit A1 toxic astrocyte profiles and conversely that SOD1^G93A^:Cx43 KO astrocytes which were neuroprotective did not show an A2 phenotype. This suggests that in the context of the SOD1^G93A^ and the SOD1^G93A^:Cx43 KO models, the toxicity of SOD1^G93A^ astrocytes and the neuroprotective effects seen with the knockdown of Cx43 appears distinct from the cytokine-activated A1/A2 phenotype profile.

Cx43 is incorporated into both GJ as well as HC. Hence, we wanted to know whether the increases in Cx43 we observed in SOD1^G93A^ mice primarily affects Cx43 HC at the astrocyte membrane. Previous studies have relied on pharmacologic blockade with peptide blockers to implicate Cx43 HC relevance in the context of disease^11, 31^. By modifying a biotin pulldown method^33^, we were able to directly demonstrate increases in the total amount of Cx43 in its HC form. Interestingly, in the SOD1^G93A^ mice the actual proportion of Cx43 in HC was significantly higher than the percentage seen in WT. Therefore, this method provides evidence that the upregulation of Cx43 may be the result of altered HC trafficking towards or from astrocytic membranes as well.

In light of our *in vivo* observations from ALS mice models regarding the potential role of Cx43 in the temporal and spatial spread of the disease, we sought to confirm our data using human ALS tissue and biofluids. We extended our previous observations^31^ regarding increases in Cx43 from an additional unique set of ALS motor cortex and cervical spinal cord samples. We observed that the expression levels of Cx43 also parallels disease course, with more aggressive forms of ALS having higher Cx43 expression levels. These data were also reflected in our finding that Cx43 levels in the CSF correlated with the temporal course of disease progression as well as the functional level of ALS patients using a clinically-relevant scale. These findings provide early evidence that Cx43 could be translationally relevant as a prognostic biomarker of disease progression.

Human iPSC-astrocytes derived from FALS patients carrying SOD1 mutations as well as from SALS patients provide a unique *in vitro* tool to investigate astrocyte contribution to neurodegeneration in the context of human pathology. Similar to the SOD1^G93A^ mouse astrocytes, we noted increases in Cx43 expression in our library of hiPSC-A lines from both FALS and SALS patients and a gene corrected isogenic SOD1^A4V^ hiPSC-A line resulted in a reduction in Cx43 expression. Our data are particularly notable as these ALS hiPSC-A have never been in co-culture with motor neurons, and, therefore, the increases in Cx43 are not determined by neuronal signals as we have previously^36^ described in hiPSC-A/-MN co-cultures, nor have they been exposed to neuronal cell death as occurs in *in vivo* ALS astrocytes.

Using a fully humanized, spinal cord specific, co-culture platform to study ALS astrocyte/MN interactions, we were able to recapitulate previous studies^24, 27, 63–67^ showing that ALS hiPSC-A were toxic to hiPSC-MN in co-culture with several FALS and SALS lines. To identify specifically whether toxicity was mediated by the extracellular release of relevant toxic factors, we used a transwell system separating ALS hiPSC-A from hiPSC-M. This system proved valuable as toxicity from ALS hiPSC-A was again noted amongst several lines. The transwell cultures, and pharmacological blockade by the Gap19 mimetic peptide, allow us to implicate the release of toxic factors through Cx43 HC into the media, an observation supported by our demonstration not only that there are more Cx43 HC on the ALS hiPSC-A membrane but they are functionally more active as well.

To examine whether the cell-autonomous and Cx43-mediated astrocyte neurotoxicity is due to ALS mutations alone, we used a gene correction strategy in the SOD1^A4V^ mutation carrying hiPSC-A. The gene editing did not result in a complete rescue of astrocyte-mediated neurotoxicity, which paralleled the relatively higher expression of Cx43 in isogenic control cells compared to control lines. This may suggest that other factors including epigenetic changes, or as of yet unidentified risk-factor genes, may be contributing to residual astrocyte-mediated neurotoxicity. Moreover, it is notable that the remaining neurotoxicity induced by isogenic SOD1^A4V^ hiPSC-A was rescued by Gap19 treatment, suggesting that Cx43 HC contribute to this astrocyte-mediated phenomenon.

Several studies have suggested that astrocyte-mediated neurotoxicity in ALS appears to be motor neuron specific^23, 25, 26, 68–73^. Our spinal cord-specific hiPSC platform of human astrocyte/neuron co-cultures may provide a more accurate window into spinal cord pathobiology. While the majority of our hiPSC-neurons are MN, there are some populations of neurons that lack either ChAT or Isl1/2 immunostaining, but express a GABAergic profile ^36^. Our data suggest that ALS hiPSC-A HC-mediated toxicity in these human cells is not specific to motor neuron subtypes but affects these other neuronal subtypes as well. This finding would be consistent with ALS being increasingly recognized as a disorder not only of motor neurons but GABAergic and glycinergic interneurons^74–76^ as well.

One of the proposed strengths of using human iPSC is their potential use in translating fundamental observations to therapeutic opportunities^77^. Mimetic peptides like Gap19 do not have the capacity to cross the blood brain barrier and therefore their clinical use is limited. Tonabersat (SB-220453)^43^, was selected for its relative specificity as Cx43 HC blocker at low concentrations^43^ and also the ability to cross the blood brain barrier^48^. Systemic delivery of tonabersat in low concentrations in a rat retinal damage model (a model for age-related macular degeneration) improved functional outcomes by electroretinography and also prevented thinning of the retina through its capacity to block Cx43 HC^43^. Tonabersat has been shown to reduce neurogenic inflammation, and to antagonize transient neuronal hyperexcitability associated with cortical spreading depression in animal models of migraine with aura^78, 79^, where a role of gap junctions has been proposed^80^. Based upon this hypothesis and preclinical results, the oral administration of tonabersat was tested in a phase II clinical trial as a prophylaxis for migraine where it was found to be well tolerated^81, 82^. Tonabersat has also been tested on animal models of epilepsy^49, 50^ and proposed as a novel antiepileptic therapy for clinical trials^83^.

We therefore investigated the effect of tonabersat on MN survival in the context of ALS. We found that tonabersat displayed dose-dependent neuroprotective effects in the ALS hiPSC-A/MN co-culture platform and even more importantly in the transwell co-culture model, consistent with its capacity to reduce Cx43 HC-mediated release of factor(s) into the media.

Since tonabersat has been shown to reduce astrocyte-mediated neuronal hyperexcitability, we used our recently established multielectrode array recording technique^36^ to test the effects of tonabersat on hiPSC-MN firing. We found that tonabersat reduces neuronal spiking and bursting activity in a dose-dependent fashion in control hiPSC-A/MN co-culture. This effect was astrocyte-specific, as the drug did not affect hiPSC-MN survival alone, and Cx43 HC-specific, as it occurred at concentrations of tonabersat known to specifically target hemichannels and was comparable to the electrophysiological effects of Gap19. We then translated this MEA platform for electrophysiological recording and drug testing to cultures of control and ALS hiPSC-A with control hiPSC-MN. We found that the culture of human motor neurons with ALS astrocytes resulted in a transient increase in neuronal firing early during the co-culture period, and, importantly, that this effect could be inhibited with acute addition of tonabersat. This study suggests, in an all human iPSC model, that tonabersat can bring control MN firing rates back to normal in co-culture with ALS astrocytes. Additional findings also support that treatment of ALS hiPSC-A/MN co-cultures with tonabersat during a defined period of neuronal hyperexcitability was sufficient to provide long-lasting neuroprotection, suggesting that there is a temporal timeframe to the induction of astrocyte-mediated MN loss.

## Conclusion

The utilization of *in vivo* and *in vitro* murine and human modeling in both FALS and SALS highlights a novel mechanism by which Cx43 HC are the conduits for ALS astrocyte-mediated MN toxicity that has a distinct temporal course, providing a potential therapeutic target for neuroprotection during the course of disease.

## Methods

### Cx43fl/fl:SOD1^G93A^:GFAP-Cre transgenic mice

All procedures were performed in accordance with the NIH Guidelines on the care and use of vertebrate animals and approved by the Institutional Animal Care and Use Committee of the Research Institute at Johns Hopkins University. Animals were housed under light:dark (12:12 h) cycle and provided with food and water *ad libitum*. Transgenic SOD1^G93A^: hGFAP-Cre: Cx43^fl/fl^ mice were generated by breeding hGFAP-Cre (Jackson labs) with Cx43^fl/fl^ mice (Jackson labs) and those mice were further crossed with B6SJ/L SOD1^G93A^ Gur/J mice (Jackson labs). The genotypes of the mice were determined by PCR using specific primer sets against each of the transgenic mice.

### Behavioral assessments in SOD1^G93A^ mice

Hindlimb and forelimb grip strength, weight, disease onset (assessed individually for each mouse by a 10% loss in hindlimb grip strength relative to each animal’s own baseline), disease duration (measured as time between hindlimb disease onset and disease endstage), and survival was conducted on SOD1^G93A^ Cx43^fl/fl^:GFAP-Cre mice, SOD1^G93A^:Cx43^fl/fl^ and SOD1^G93A^:GFAP-Cre mice^84^.

### In vivo immunohistochemistry

To examine the pathology in transgenic SOD1^G93A^ mice, we sacrificed mice at several time points. Mice were anesthetized and trans-cardially perfused first with saline and then with 4% PFA. Brain and spinal cord were isolated and fixed overnight in 4% PFA. The tissue was rinsed with 0.2 M PB the next day and cryopreserved in 30% sucrose solution. The tissue was then frozen in tissue freezing medium and sectioned at 20 μm thickness. For immunohistochemistry, in brief, spinal cord sections were rinsed three times with 0.1 M PBS for 10 min each. The sections were then blocked with 10% goat block containing 0.2% triton-X for 1 hour. They were incubated overnight at 4°C with the relevant antibodies in 5% goat block. The next day the sections were washed in 0.1M PBS and then incubated with the species-specific secondary antibody for two hours at room temperature. The sections were then washed and mounted with Prolong gold with DAPI (Life Technologies) and stored until ready to image as we have described previously^31^. The antibodies used are detailed in **Table 3.**

**Table 3.** Primary antibodies used for in vitro and in vivo immunostaining.

| Antibody | Species | Dilution | Company (cat no) |
| --- | --- | --- | --- |
| Cx43 | Rabbit | 1:500 | Millipore Sigma (C6219) |
| GFAP | Chicken | 1:200 | Millipore sigma (AB5541) |
| GFAP | Mouse | 1:200 | Millipore Sigma (MAB360) |
| Tuj1 | Rabbit | 1:500 | Millipore Sigma (AB9354) |
| ChAT | Goat | 1:100 | Millipore Sigma (AB144P) |
| Isl1/2 | Mouse | 1:50 | Developmental Studies Hybridoma Bank (39.4D5, concentrate) |
| Iba-1 | Rabbit | 1:100 | Wako (019-19741) |
| GLT-1 | Rabbit | 1:500 | (*) |

(\*) Kindly provided by Dr Jeffrey D. Rothstein's Lab

### Tissue immunoblot (Western Blot)

Aliquots of homogenized samples from mouse spinal cord regions were separately subjected to SDS-polyacrylamide gel electrophoresis. Blots were probed with primary antibodies: rabbit polyclonal Connexin 43 (1:5000, Sigma-Aldrich, C6219) and mouse monoclonal GFAP (1:5000, Millipore Sigma, MAB360). Immunoreactivity was visualized by enhanced chemiluminescence, quantified densitometrically using Image J software, and normalized to the loading control (GAPDH). Western Blot for human tissue was performed similarly.

### In vivo motor neuron survival

The spinal cord was serially sectioned transversely (20 µm) and stained with SMI-32 to quantify motor neurons. Only motor neurons with a clearly identifiable nucleus and nucleolus, a cell soma over 400 µm^2^ and located within the ventral horn were counted and represented unilaterally^85^.

### Murine astrocytes culture

Glial restricted precursors (GRPs) were isolated from embryonic mouse spinal cord at 11.5 days and cultured as previously described^86, 87^. GRPs were then differentiated into astrocytes by supplementing the media with 10% FBS and then plated in confluent monolayers for one week prior to use.

### Murine astrocyte/motor neuron co-culture

Astrocyte motor neuron co-culture experiments were performed as described previously by Haidet-Phillips et al^88, 89^. Briefly, mouse ES cells that express GFP driven by the Hb9 promoter were differentiated to MN as previously described^90^. Three days prior to MN cell sorting, astrocytes from SOD1^G93A^ mice were plated on 96 well plates. FACS purified Hb9-GFP+ MN were plated onto astrocytes at 10,000 cells/well in MN medium^26^and were assessed by taking daily images of the GFP expressing MN to quantify the number of MN surviving per well and were normalized to day one counts.

### In vitro immunocytochemistry

Mouse astrocyte-motor neuron co-cultures were fixed with 4% PFA for 10 min, then finally blocked with 5% fetal bovine serum (FBS) for 1 hour. The cells were then stained with the appropriate primary and secondary antibodies as detailed in **Table 3**. For quantification purposes, 5-10 images were obtained from each coverslip and at least 3 coverslips were utilized for each condition. Images were analyzed using Zen 2 software (Zeiss), blinded to experimental conditions. All immunofluorescence images were arranged using Adobe Photoshop. Immunocytochemistry on coverslips from hiPSC-A/-MN co-cultures was performed using the same protocol.

### qPCR of A1/A2 transcripts

Rodent astrocytes were plated in six well plates at a density of 100,000 cells/well. Total RNA was extracted using RNeasy Mini Kit (Qiagen, 74104) and then treated with DNase I (Qiagen, 79254) to remove DNA contamination according to manufacturer’s instructions. The RNA concentration was then measured using a NanoDrop spectrophotometer (Thermo Fisher Scientific) and all RIN values were in the 2.0-2.5 range. One microgram of RNA was converted to cDNA using the iScriptTM cDNA synthesis kit (Bio-Rad). Primers for A1/A2 transcripts were ordered directly from Liddelow et al^28^. After testing primers for successful melting curves and Ct values in the appropriate range, samples were run at 1:5 or 1:10 dilution with a working concentration of 10 uM primers and Fast SYBR green master mix (Applied Biosystems, 4385610). Ct values were transformed to ΔΔCt values by normalizing to 18s and WT samples. 18s primers were run on each plate to account for technical variability.

### Cx43 biotinylation assay

The biotin pulldown assay was performed as previously described^33^ and adapted for use in our disease models. The biotin binds to lysine residues on the extracellular side of membrane proteins in intact cells. The lysine residue of Cx43 is accessible to binding when in HC but not in GJ formation, which ensures the specificity of HC biotin tagging.

Rodent astrocytes were plated in 6 well plates at a density of 500,000 cells/well and live cells were treated with EZ-LinkTMNHS-Biotin reagent (Thermo Scientific, 20217). Cells were then homogenized and pelleted. Streptavidin beads (Pierce^TM^ Streptavidin Agarose, 20349) were then added, the mixture was pelleted, washed and then heated to remove the streptavidin from the protein of interest. The samples were finally suspended in sample buffer and frozen at −80°C or used immediately for immunoblot. For analysis, intensities of Cx43 total protein, ranging from one to 20µg of loaded protein, were compared densitometrically to form a linear scale. On the same blot, triplicate wells of the biotin pulldown fraction were then measured densitometrically and mapped onto the scale, with separate linear plots for WT and SOD1^G93A^ samples. Once the densities of the HC were measured and located on the linear curve, the corresponding µg amount on the x axis was then compared to the total amount of initially collected protein. The amount of HC was then converted to a percentage of total Cx43 protein. Linear scales were only accepted with R^2^ > 0.95. Calculations include correction for 10% loading of biotin pulldown samples in order to allow for intensity readings to fit on the total protein intensity scale^33^. For human iPSC-A, the cells were grown according to the below protocol and protein was extracted in the same manner as described for rodent astrocytes. For the total protein scales, the range was decreased from one to 12µg to accommodate increased staining intensities in the protein.

### Human tissue

Human samples from control and sporadic ALS patients were obtained from the “VA Tissue Bank” and the “Target ALS - Multicenter Human Postmortem Tissue Core*”* (http://www.targetals.org/human-postmortem-tissue-core/). The samples utilized in the study are described in **Table 1**. Western blot on human tissue was conducted using similar protocol to mouse tissues described above.

### Cerebrospinal fluid measurements of Cx43

Cerebrospinal fluid (CSF) was obtained from the NEALS Biorepository (https://www.neals.org/for-als-researchers/neals-sample-repository/) including 8 control and 15 ALS patient samples. The degree of ALS disease progression was categorized^34, 35^ as slow (ALSFRS-R <0.5 points/month), typical (>0.5 and <1.0 points/month), or rapid progression (>1.0 point/month) at the time of sample collection. Healthy control samples were also included in the analysis. ELISA was performed according to the manufacturer’s protocol, specifically designed for detection of Cx43 in human cells (BioMatik, EKU03444). In short, CSF samples were taken directly from the NEALS aliquots and added to each well of the pre-antibody-bound plates in technical triplicate, in addition to a Cx43 standard ranging from 10ng/mL to 0.156 ng/mL and a blank. After the addition of the detection reagents, the wells were thoroughly washed before adding the substrate and stop solutions. The plates were then immediately read at 450nm using a microplate reader (SpectraMax M3, Molecular Devices).

### Human iPSC astrocyte culture

The generation, characterization of the human iPSC lines and their differentiation into astrocytes have been previously described by our group^26, 36, 91^. Briefly, our 25-30 day spinal cord differentiation protocol relies on extrinsic morphogenetic instructions to pattern neural progenitor cells (NPC) along the rostro-caudal and dorso-ventral axis. For the generation of hiPSC-A, NPC were cultured for up to 90 days in vitro (DIV) with “astrocyte differentiating medium”^36^. The human iPSC-derived astrocyte lines from healthy subjects and both sporadic and familial ALS patients utilized for this study are listed in **Table 2**. For Western Blot purposes, hiPSC-A from healthy subjects and ALS patients were cultured for 86 DIV and then plated in 6 well plates at a density of 200,000 cells/wells. Astrocytes were allowed to recover and become confluent for 4 days (i.e. 90 DIV) prior to collection of protein samples and immunoblotting.

### ATP detection

In order to detect ATP in the supernatant and from total cells, hiPSC-A were cultured according to the above protocol. After astrocytes were matured past the 90 day mark, cells were plated at 15,000 cells/well in a 96 well plate for one week. After one week of being plated, cells were treated with either vehicle (fresh media) or Gap19 (Tocris, 53533) at a concentration^31, 36^ of 340 µM. Cells were kept in 150µL of media which was replaced 50% each feed with either vehicle or Gap19 every other day. Nine days after initial treatment, 75 µL of supernatant from each well was collected for ATP supernatant detection using the ATPlite Luminescence Assay Kit (Perkin Elmer, 6016943) as described by the kit protocol. Detection was conducted on white-opaque plates using the Perkin Elmer EnVision 2105 Multimode Plate Reader (Perkin Elmer 2015-0010) using the Ultra-Sensitive Luminescence Detection mode. Total cell measurements were then conducted by collecting the cells in the kit provided lysis buffer and measuring their luminescence post luciferase treatment according to the kit instructions on white-opaque plates again using the EnVision Plate Reader.

### Human iPSC-motor neuron culture

Briefly, NPC differentiated with the aforementioned spinal cord patterning protocol were cultured for up to 60 DIV with “neuronal differentiating medium”^36, 92^. To prevent astrocyte over-proliferation, neuronal cultures were treated once with 0.02µM cytosine arabinoside (Ara-C) (Millipore Sigma) for 48 h. While the majority of these cells are Tuj1^+^/ChAT^+^, consistent with a spinal motor neuronal phenotype, a smaller subset of neurons (Tuj1^+^/ChAT^-^) is also present and consists mainly of GABAergic interneurons^36^.

### Human iPSC-astrocyte and hiPSC-motor neuron co-culture

Human iPSC-A form healthy subjects and ALS patients were cultured for 86 DIV and then plated in 24 well plates on glass cover-slips at a density of 20,000 cells/well. Four days later, hiPSC-MN from control samples, cultured for 60 DIV prior to use, were plated on the top of astrocytes at a density of 50,000 cells/well. This was considered day 1 of co-culture. Co-cultures were maintained in motor neuronal medium enriched with 1% FBS for a total of 14 days before fixing for immunocytochemistry. Co-culture medium was changed every third day with half medium exchange.

For pharmacological assays on Cx43 HC, we used GAP19 (Tocris, 53533) at a concentration of 340 µM ^31, 36^ and Tonabersat (SB-220453) at concentrations of 1 µM, 10 µM and 100 µM ^43, 44^ (Millipore Sigma, SML1354).

### Human iPSC-astrocyte and hiPSC-motor neuron co-culture in a transwell system

Human iPSC-MN from control samples were cultured for 56 DIV and then plated in 12 well plates on glass coverslips at a density of 250,000 cells/well. In parallel cultures, hiPSC-A from healthy subjects and ALS patients were cultured for 86 DIV and then plated in the top inserts of 12 well plates (12 mm diameter, with 0.4µm pore polyester membrane insert) (Corning, 3460) at a density of 48,000 per insert. Four days later, the inserts with 90 DIV astrocytes were placed on the top of 12-well plates with 60 DIV motor neurons at the bottom. This was considered day 1 of “transwell” co-cultures. Half medium exchanges were performed from both the top and the bottom compartments. Transwell co-culture were otherwise maintained as conventional co-cultures and neurons on glass coverslips were fixed for immunocytochemistry after 14 days.

### Isogenic A4V line generation and Sanger sequencing

The SOD1^A4V/+^ line and its corrected isogenic control (SOD1^+/+^) have been previously utilized in a published^38^ study by one of the authors (KE). The SOD1 A4V mutation has been corrected with zinc-finger nuclease editing. In this study, we confirmed gene correction in the SOD1^+/+^ pluripotent stem cells, and verified that it was retained later during differentiation in neural progenitor cells and astrocytes (hiPSC-A). Briefly, genomic DNA was extracted using QIAamp UCP DNA Micro Kit (Qiagen, 56204) and expanded with REPLI midi Kit (Qiagen, 150043). SOD1 Exon 1 was expanded using 20 µM of forward (CTATAAAGTAGTCGCGGAGACGGGGTG) and reverse (GCGGCCTCGCAAACAAGCCT) primers with Phusion Hot Start II DNA Polymerase (Thermo Fisher, F549L), identified and isolated in 1% agarose gel, and then sequenced by Genewiz©.

### Multi-electrode array culture and recordings

Multi-electrode array (MEA) plates (MultiChannel Systems, 60MEA200/30iR-Ti-gr,) with 60 electrodes, including 59 active and 1 reference, and a MEA2100 (MultiChannel Systems) platform were used for the recording of neuronal activity in hiPSC-A/-MN co-cultures as we have previously described^36^. Briefly, hiPSC-A from healthy subjects and ALS patients and hiPSC-MN from a single control line were serially co-cultured at densities of 100,000 cells/plate and 500,000 cells/plate, respectively. We analyzed the following electrophysiological parameters: spike rate, burst rate, and percentage of spiking and bursting electrodes (on the overall 59 recording electrodes). The recording of the baseline activity of MEA plates was performed every 3 days over 1-minute period for 2 weeks after initial plating. For pharmacological assays, we performed 5 minutes recordings to capture slower electrophysiological effects of HC blockers^36^: baseline MEA activity for 1 minute, activity for 1 minute after 100 µl medium exchange with drug vehicle, and activity for 3 minutes after the exchange of 100 µl of medium containing the compound of interest and the drug vehicle. For analysis purposes, we considered only the electrodes with spike rate > 0. Spiking and bursting activity is expressed as the mean of individual MEA plates.

### Data presentation and statistical analysis

All data were analyzed using GraphPad Prism software (La Jolla, CA). Graph bars represent mean ± SEM and individual observations are shown as scatter plots. Data were analyzed using Student’s t-test, one-way ANOVA or two-way ANOVA, followed by Tukey’s test for multiple comparisons as described in figure legends. We did not perform a sample-size calculation, but sample-sizes are similar to those reported in the literature, including work from our laboratory and co-authors laboratories, as well. We assumed that the data distribution was normal but we did not perform any normality distribution test. The statistical significance was set at p < 0.05 (*p < 0.05, **p < 0.01, ***p < 0.001). All experiments were performed in at least 3 technical and/or biological replicates. The number (n) of experimental conditions is indicated in the figure legends.

## Acknowledgements

ALS Association 18-DDC-436 (NJM), Maryland Stem Cell Research Fund 2019-MSCRFD-5122 (NJM), Packard Center for ALS research at Johns Hopkins (NJM), Department of Defense ALSRP W81XWH1810175 (NJM). NIH R01-GM099490 (JEC). The SOD1^A4V^ isogenic lines were a kind gift from Kevin Eggan. ALS and control brain and spinal cord tissues were obtained from the Target ALS Human Postmortem Tissue Core (Lyle Ostrow) as well as the VA Tissue Bank. Northeast ALS Consortium Sample Repository provided CSF samples (James Berry). Illustrations were created with BioRender (https://biorender.com/).

## Author Contributions

Akshata Almad: Conception and design of work, acquisition and analysis of data, interpretation of data, manuscript draft and revision.

Arens Taga: Conception and design of work, acquisition and analysis of data, interpretation of data, manuscript draft and revision.

Jessica Joseph: Conception and design of work, acquisition and analysis of data, interpretation of data, manuscript draft and revision.

Connor Welsh: Acquisition and analysis of data

Aneesh Patankar: Acquisition and analysis of data

Sarah Gross: Acquisition and analysis of data, interpretation of data, manuscript draft and revision.

Jean-Philippe Richard: Acquisition and analysis of data, interpretation of data, manuscript draft and revision.

Aayush Pokharel: Acquisition and analysis of data

Mauricio Lillo: Acquisition and analysis of data, interpretation of data, manuscript draft and revision.

Raha Dastgheyb: Acquisition and analysis of data, interpretation of data, manuscript draft and revision.

Kevin Eggan: Interpretation of data, manuscript draft and revision.

Norman Haughey: Acquisition and analysis of data, interpretation of data, manuscript draft and revision.

Jorge Contreras: Conception and design of work, acquisition and analysis of data, interpretation of data, manuscript draft and revision.

Nicholas J. Maragakis: Conception and design of work, acquisition and analysis of data, interpretation of data, manuscript draft and revision.

## Dataset Availability

The datasets generated during and analyzed during the current study are available from the corresponding author on reasonable request.

## Ethics Statement

The protocols for the use of human iPSC have undergone and received Johns Hopkins University IRB approval. The titles of the protocols are “Skin biopsies and peripheral blood mononuclear cells (PBMCs) generate cell lines for study of Amyotrophic Lateral Sclerosis” (approval 10/16/2008--application number NA_00021979) and “Generation and characterization of cell lines for Amyotrophic Lateral Sclerosis” (approval 1/19/2010--application number NA_00033726). These protocols are currently active. Informed consent was obtained from all donors. Donations of samples was voluntary.

---

**Supplementary Figure 1. Cx30 is elevated in SOD1^G93A^:Cx43 KO mice, with no change in GLT-1 and Iba-1. (a)** The expression of Cx30 was significantly elevated in the lumbar ventral grey matter (VGM) of and SOD1^G93A^:Cx43 KO mice compared to control SOD1 mice at endstage (t-test, ***p<0.001). **(b)** No changes were observed in Iba1 immunoreactivity in the VGM at the endstage. **(c, d)** Similarly no changes were observed in GLT1 based on immunoreactivity and immunoblotting. n=3-4 animals/group, and at least 3 sections /animal for immunohistochemistry.

**Supplementary Figure 2. Cx43 deletion in SOD1^G93A^ astrocytes increases survival of mouse motor neurons. (a,b)** Astrocytes were from SOD1^WT^, SOD1^G93A^ and SOD1^G93A^:Cx43KO mice, and while Cx43 increases in SOD1^G93A^ astrocytes, a dramatic loss of Cx43 was confirmed in SOD1^G93A^:Cx43KO astrocytes using IHC and immunoblotting analysis. One-way Anova, *p<0.05, n=3. **(c)** Hb9-GFP expressing motor neurons were plated on top of SOD1^WT^, SOD1^G93A^ and SOD1^G93A^:Cx43KO astrocytes. While motor neurons plated on SOD1^G93A^ astrocytes degenerated rapidly, significant neuroprotection was observed when motor neurons were plated on SOD1^G93A^ Cx43KO astrocytes. One-way Anova, ***p<0.001, n=3.

**Supplementary Figure 3. A1 and A2 transcripts in SOD1^G93A^ and SOD1^G93A^:Cx43KO astrocytes. (a)** Gene profile of activated astrocytes does not show increases in A1 toxic transcripts in SOD1^G93A^ astrocytes nor does the KO of Cx43 result in a reduction of A1 transcripts. **(b)** A2 transcript quantification does not show a specific pattern of expression in SOD1^G93A^ or SOD1^G93A^:Cx43KO astrocytes. Bars are mean values of n=3 technical replicates.

**Supplementary Figure 4. Spinal cord directed hiPSC-A and hiPSC-MN co-culture. (a)** Schematic representation of hiPSC differentiation into spinal cord-specific hiPSC-A and MN. **(b)** Representative immunostaining of hiPSC-A for GFAP (green) and hiPSC-MN for TUJ1 in co-culture. **(c)** Human iPSC-MN immunostained for choline acetyltransferase (ChAT) (green). A ChAT^-^ neuron is marked (arrow) as well. **(d)** A subset of hiPSC-MN immunostained for the MN marker Isl1 (green). An Isl1^-^ is marked (arrow).

**Supplementary Figure 5. Gene correction of SOD1^A4V^ in hiPSC-A results in reduced Cx43 expression, and residual non-SOD1-mediated MN toxicity which is partially rescued by Gap19. (a)** Schematic representation of SOD1 gene A4V correction. **(b)** The SOD1^A4V^ hiPSC-A line (39B SOD1^A4V/+^) shows an increase in Cx43 expression that is partially normalized by gene correction seen in line 39B2.5(SOD1^+/+^). The residual Cx43 expression in the isogenic (SOD1^+/+^) line is significantly higher than control line. One-way Anova, * p<0.05; ^^^ p<0.001, n=3 **(c-m)** The isogenic gene-corrected line 392.5B (SOD1^+/+^) shows less toxicity to ChAT^+^ and Isl1^+^ hiPSC-MN when compared to uncorrected line SOD1^A4V/+^ in a co-culture paradigm. In both lines, however, additional neuroprotection was obtained following addition of Gap19. **(n-x)** Transwell co-culture of SOD1^A4V/+^ astrocytes with hiPSC-MN shows expected neurotoxicity which is less evident in the SOD1^+/+^ hiPSC-A/control MN co-culture. Additional neuroprotection was also evident following Cx43 HC specific blockade with Gap19. One-way ANOVA, * or ^ p<0.05; ** or ^^ p<0.01; *** or ^^^ p<0.001, n=3.

**Supplementary Figure 6. Blocking Cx43 HC with Gap19 has no effect on control hiPSC-MN cultured either with control hiPSC-A or when cultured alone. (a)** The transwell co-culture of control hiPSC-A/control MN or control hiPSC-MN alone. **(b)** The application of Gap19 has no effect on ChAT+ and Isl1+ control hiPSC-MN survival, which was indipendent of culture conditions (alone or in co-culture with control hiPSC-A). One-way ANOVA. One-way ANOVA, n=3.

**Supplementary Figure 7. Cx43 HC blockade results in neuroprotection of ALS hiPSC-A induced non-motor neuron cell death. (a)** FALS and SALS hiPSC-A induce hiPSC non-motor neuron cell death in a co-culture system as defined by ChAT^-^/Tuj1^+^ and Isl1^-^/Tuj1^+^ immunoreactivity that can be rescued with Gap19. (**b)** The transwell culture of FALS and SALS hiPSC-A induce non-motor neuron cell death which can be rescued with Gap19. One-way ANOVA, * or ^ p<0.05; ** or ^^ p<0.01; *** or ^^^ p<0.001, n=3.

**Supplementary Figure 8. ALS hiPSC-A toxicity towards motor and non-motor neurons and Cx43 HC-mediated neuroprotection was evaluated in an additional control hiPSC line (CS25) using a transwell culture. (a)** FALS and SALS hiPSC-A induce toxicity in hiPSC-motor neuronal populations (ChAT+/Tuj1+ and Isl+/Tuj1+) differentediated from the control CS25 hiPSC-line. Motor neuronal death is partially ameliorated by the addition of Gap19, a specific Cx43 HC blocker. **(b)** FALS and SALS hiPSC-A induce cell death in hiPSC non-motor neurons, as defined by ChAT-/Tuj1+ and Isl1-/Tuj1+ immunoreactivity, differentiated from the CS25 control hiPSC-line. Non-motor neuronal death is partially rescued by Cx43 HC blockade with Gap19. One-way ANOVA, * or ^ p<0.05; ** or ^^ p<0.01; *** or ^^^ p<0.001, n=3.

**Supplementary Figure 9. Tonabersat results in neuroprotection of ALS hiPSC-A induced non-motor neuron cell death. (a)** FALS and SALS hiPSC-A induce hiPSC non-motor neuron cell death in a co-culture system as defined by as defined by ChAT^-^/Tuj1^+^ and Isl1^-^/Tuj1^+^ immunoreactivity that can be rescued with tonabersat. **(b)** The transwell culture of FALS and SALS hiPSC-A induce non-motor neuron cell death which can be rescued with tonabersat. One-way ANOVA, * or ^ p<0.05; ** or ^^ p<0.01; *** or ^^^ p<0.001, n=3.

**Supplementary Figure 10. Tonabersat has no effect on control hiPSC-MN cultured either with control hiPSC-A or when cultured alone. (a)** The transwell co-culture of control hiPSC-A/control MN or control hiPSC-MN alone. **(b)** The application of tonabersat at several concentrations has no effect on ChAT+ and Isl1+ control hiPSC-MN survival. One-way ANOVA, n=3.

**Supplementary Figure 11. Multielectrode array recordings from hiPSC-A/MN co-cultures. (a)** Control hiPSC-MN and hiPSC-A were plated on multielectrode array (MEA) plates for recording electrophysiological activity (spike rate, percentage of overall electrodes displaying spiking or bursting activity). **(b)** MEA electrophysiological activity in control hiPSC-A/-MN co-cultures 5 minutes after the application of vehicle (-), 1µM (+) and 10 µM (++) tonabersat. **(c)** MEA electrophysiological activity in control hiPSC-A/-MN co-cultures 5 minutes after the application of vehicle (-) or 340 µM Gap19 (+). **(d)** The addition of 10 µM (++) tonabersat to cultures of control hiPSC alone does not result in an alteration of electrophysiological activity. The electrophysiological parameters are normalized to the baseline activity recorded for 1 minute (dashed line). Data are the mean of n=3 independent experiments per drug. One-way ANOVA, * p<0.05; ** p<0.01; *** p<0.001.

