## Supplemental figures for "Connexin 43 hemichannels mediate spatial and temporal disease spread in ALS"

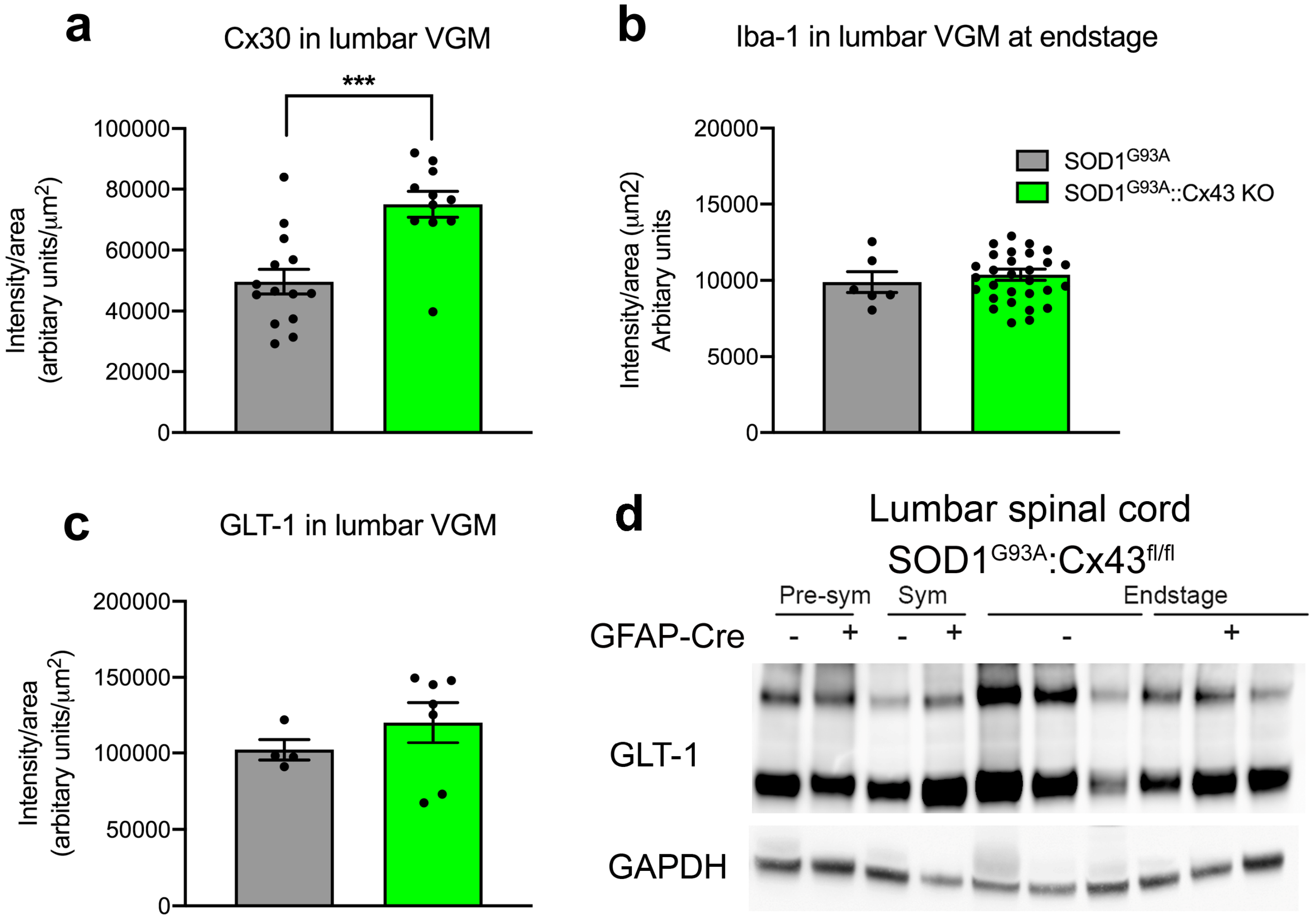
a
Cx30 in lumbar VGM
***
Intensity/area (arbitary units/μm2)
100000
80000
60000
40000
20000
0
b
Iba-1 in lumbar VGM at endstage
Intensity/area (μm2) Arbitary units
20000
15000
10000
5000
0
SOD1G93A
SOD1G93A::Cx43 KO
c
GLT-1 in lumbar VGM
Intensity/area (arbitary units/μm2)
200000
150000
100000
50000
0
d
Lumbar spinal cord
SOD1G93A:Cx43fl/fl
Pre-sym
Sym
Endstage
GFAP-Cre
- + - + - +
GLT-1
GAPDH

Fig.S1

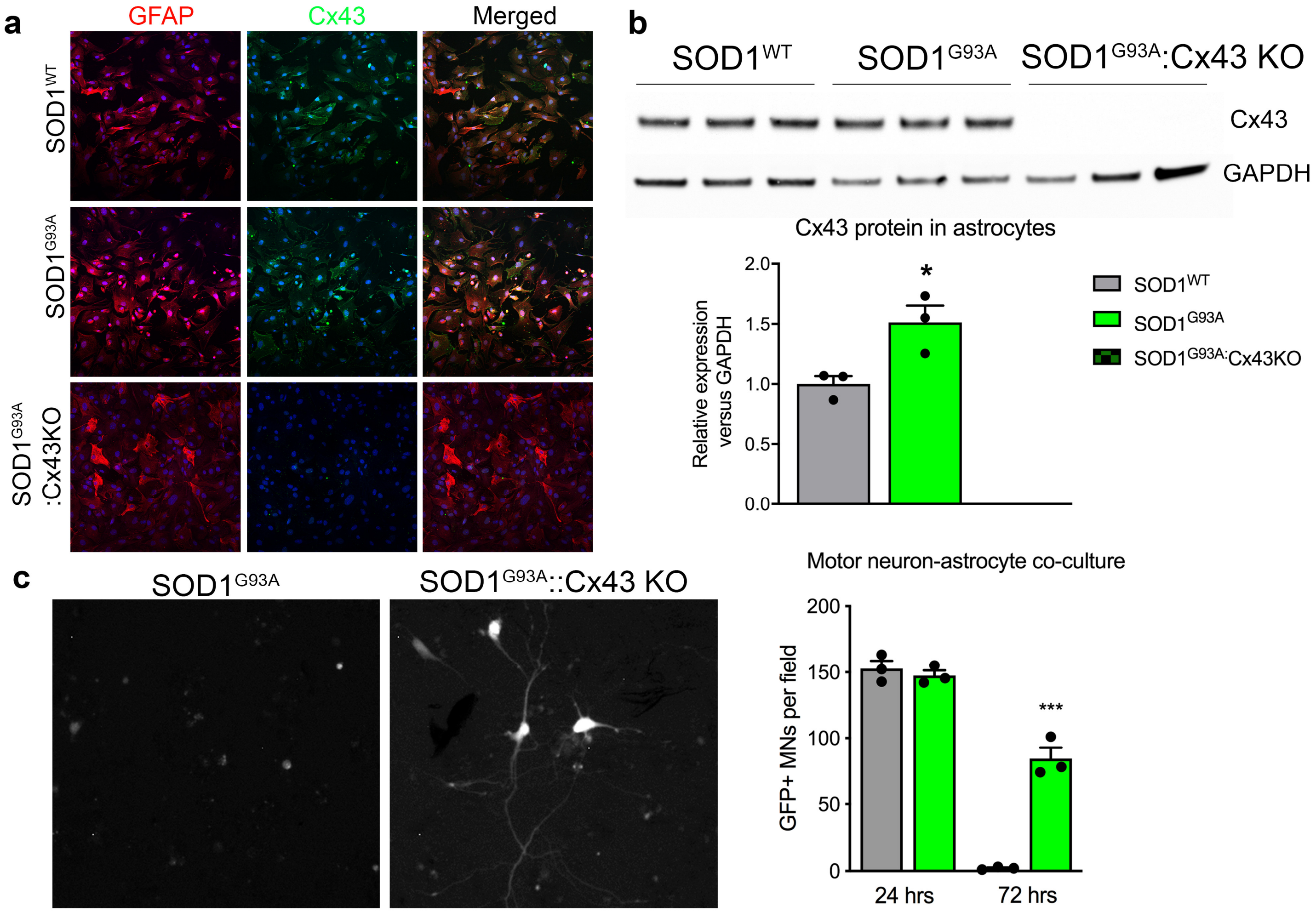

a
GFAP
Cx43
Merged
SOD1WT
SOD1G93A
SOD1G93A :Cx43KO
b
SOD1WT
SOD1G93A
SOD1G93A:Cx43 KO
Cx43
GAPDH
Cx43 protein in astrocytes
Relative expression versus GAPDH
2.0
1.5
1.0
0.5
0.0
*
SOD1WT
SOD1G93A
SOD1G93A:Cx43KO
c
SOD1G93A
SOD1G93A::Cx43 KO
Motor neuron-astrocyte co-culture
GFP+ MNs per field
200
150
100
50
0
***
24 hrs
72 hrs


Fig.S2

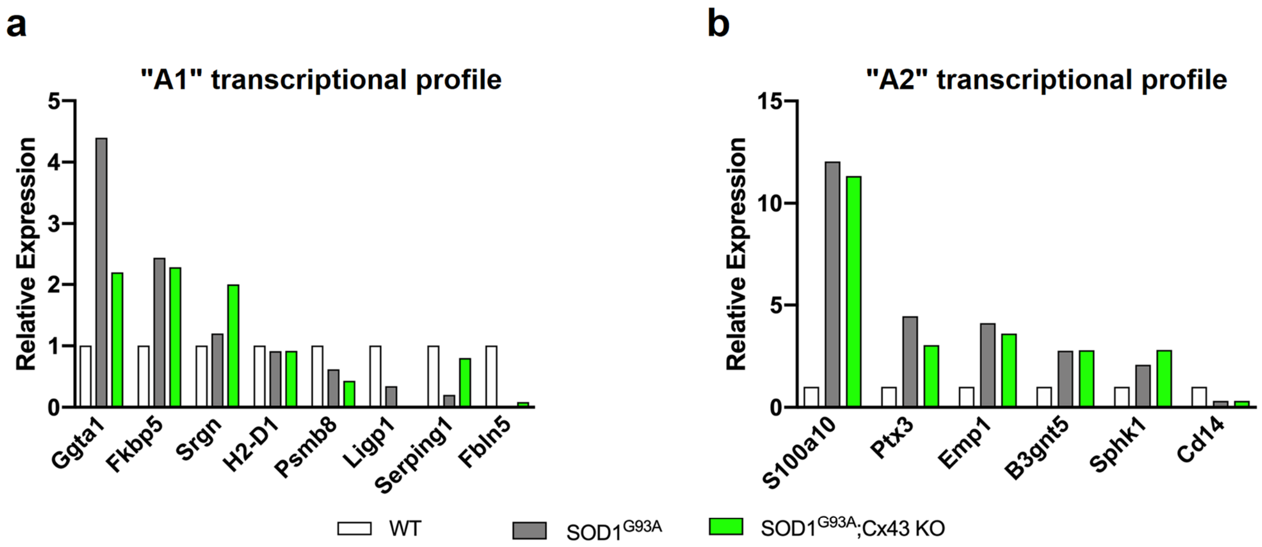
a
"A1" transcriptional profile
Relative Expression
5
4
3
2
1
0
Ggta1
Fkbp5
Srgn
H2-D1
Psmb8
Ligp1
Serping1
Fbln5
b
"A2" transcriptional profile
Relative Expression
15
10
5
0
S100a10
Ptx3
Emp1
B3gnt5
Sphk1
Cd14
WT
SOD1G93A
SOD1G93A;Cx43 KO

Fig.S3

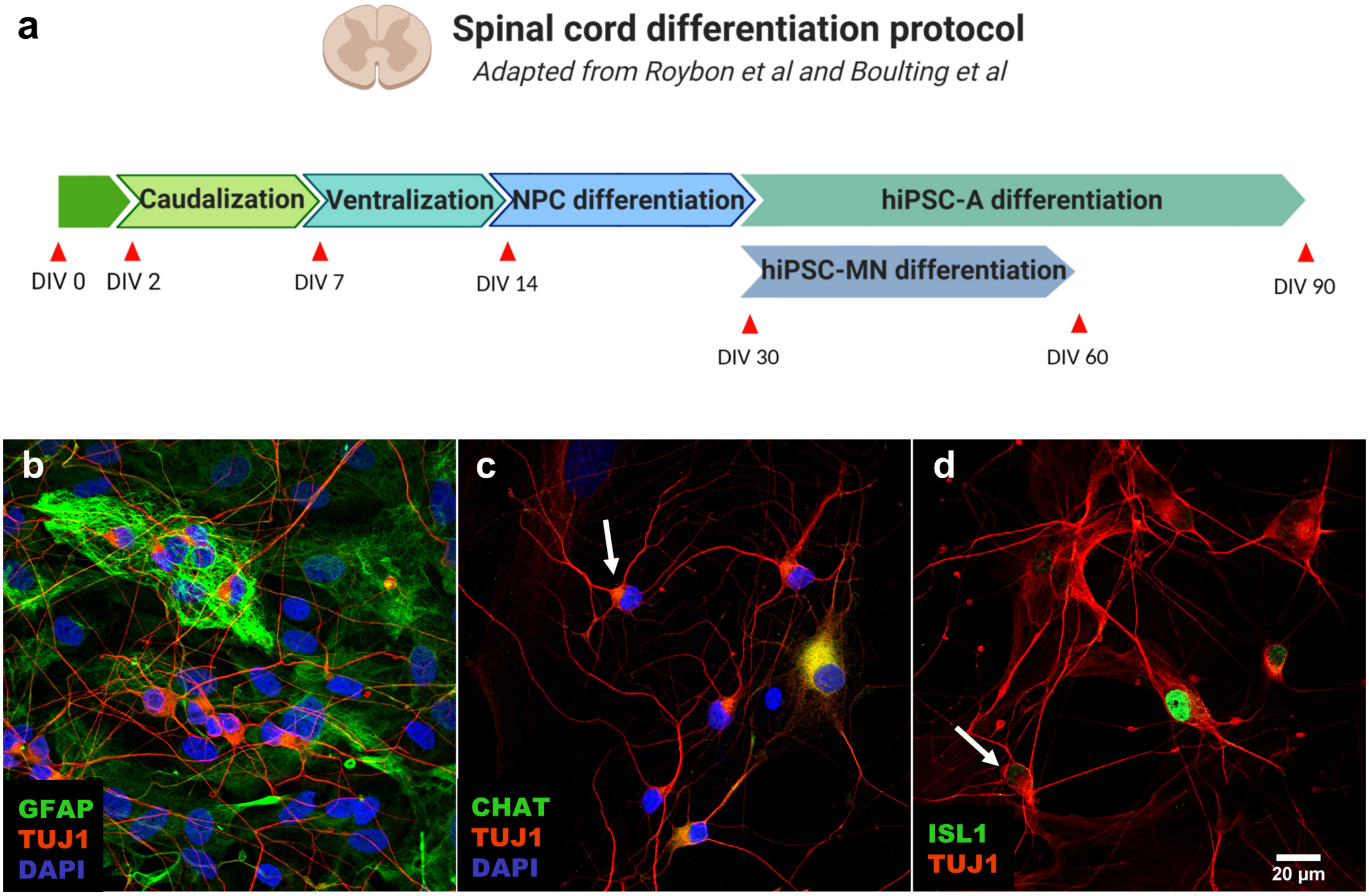
a
Spinal cord differentiation protocol
Adapted from Roybon et al and Boulting et al
Caudalization
Ventralization
NPC differentiation
hiPSC-A differentiation
hiPSC-MN differentiation
DIV 0
DIV 2
DIV 7
DIV 14
DIV 30
DIV 60
DIV 90
b
GFAP
TUJ1
DAPI
c
CHAT
TUJ1
DAPI
d
ISL1
TUJ1
20 µm

Fig.S4

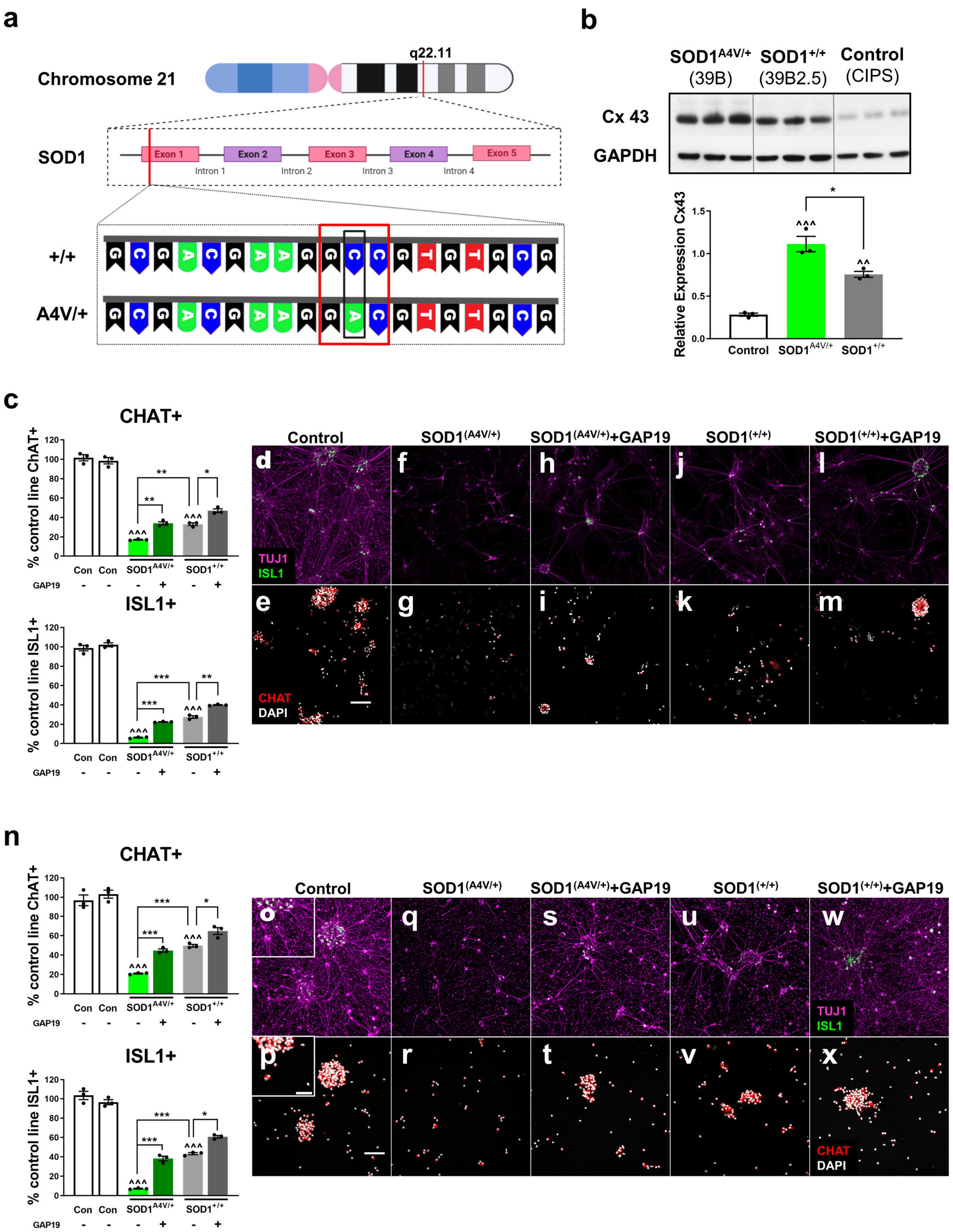

a
Chromosome 21
q22.11
SOD1
Exon 1
Exon 2
Exon 3
Exon 4
Exon 5
Intron 1
Intron 2
Intron 3
Intron 4
+/+
A4V/+
b
SOD1A4V/+ (39B)
SOD1+/+ (39B2.5)
Control (CIPS)
Cx 43
GAPDH
Relative Expression Cx43
Control
SOD1A4V/+
SOD1+/+
c
CHAT+
% control line ChAT+
ISL1+
% control line ISL1+
Con
GAP19
Control
SOD1(A4V/+)
SOD1(A4V/+)+GAP19
SOD1(+/+)
SOD1(+/+)+GAP19
TUJ1
ISL1
CHAT
DAPI
n


Fig.S5

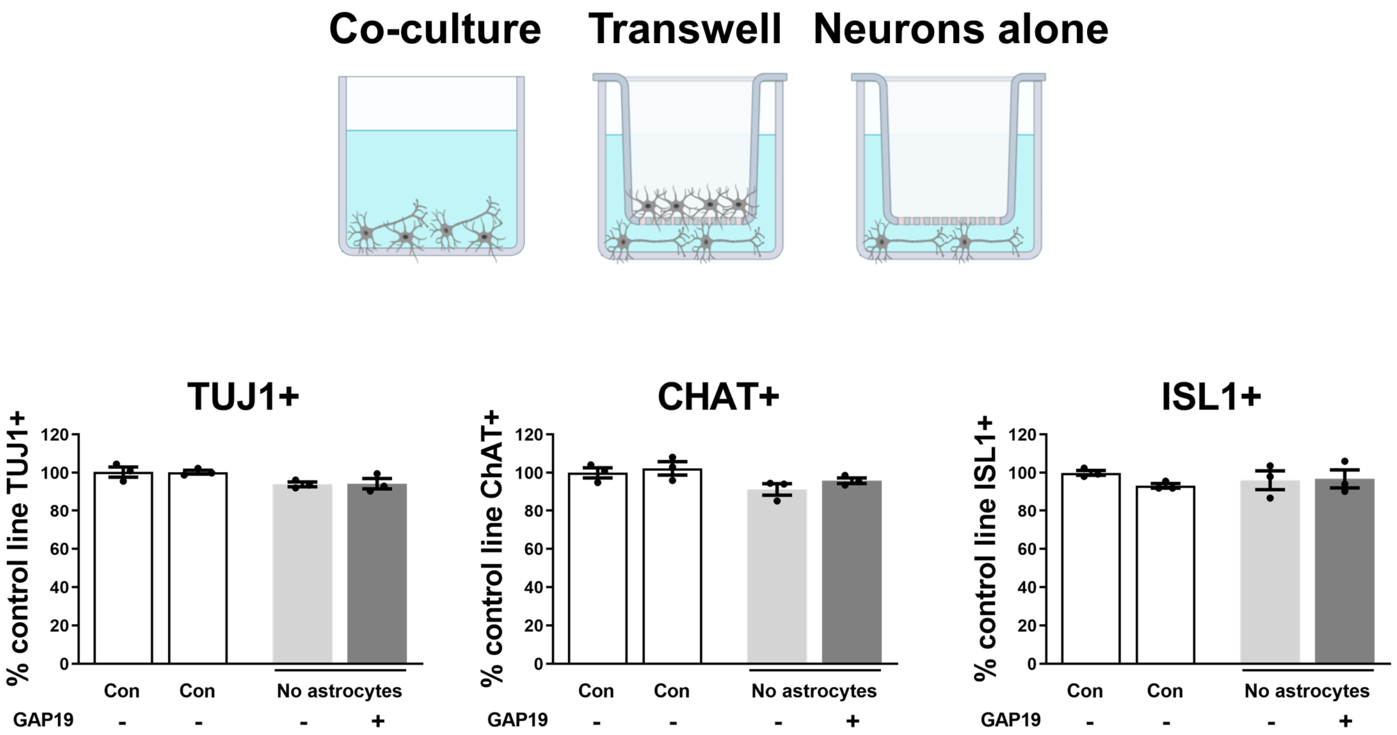
Co-culture
Transwell
Neurons alone
TUJ1+
% control line TUJ1+
CHAT+
% control line ChAT+
ISL1+
% control line ISL1+
0
20
40
60
80
100
120
Con
Con
No astrocytes
GAP19
-
-
-
+

Fig.S6

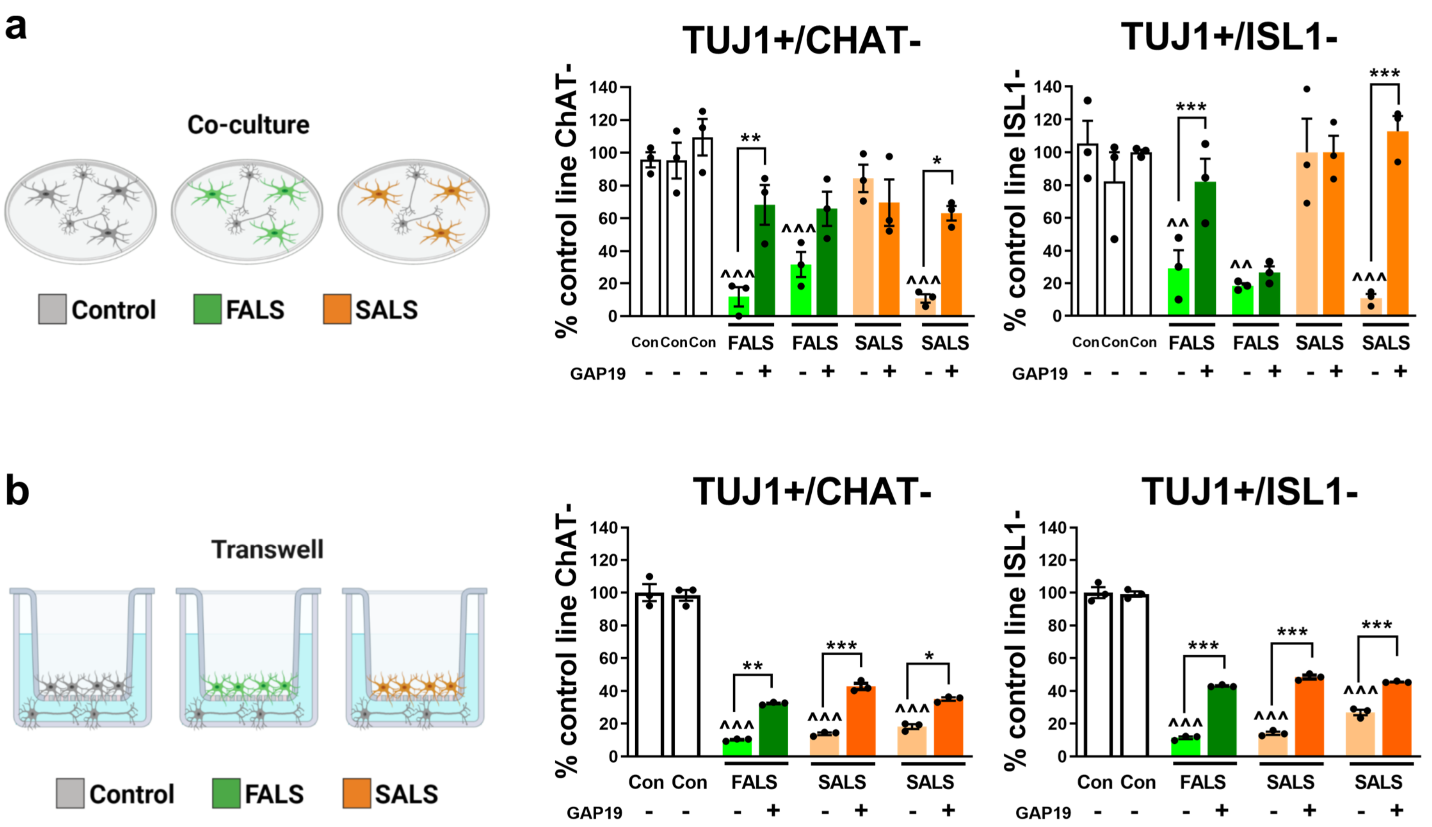
a
Co-culture
Control
FALS
SALS
TUJ1+/CHAT-
% control line ChAT-
Con Con Con FALS FALS SALS SALS
GAP19
TUJ1+/ISL1-
% control line ISL1-
b
Transwell
Control
FALS
SALS
TUJ1+/CHAT-
% control line ChAT-
Con Con FALS SALS SALS
GAP19
TUJ1+/ISL1-
% control line ISL1-

Fig.S7

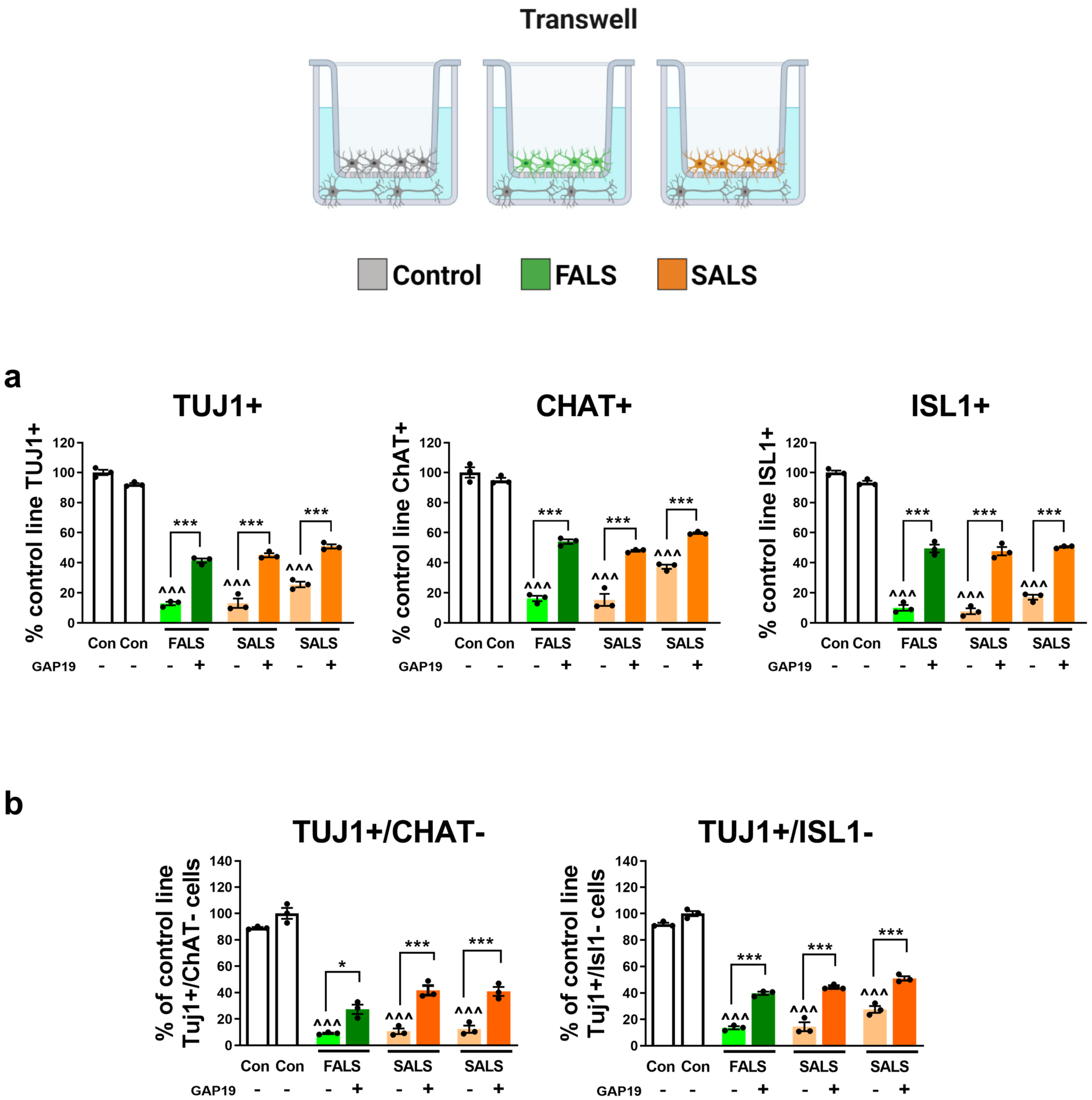
Transwell
Control
FALS
SALS
a
TUJ1+
% control line TUJ1+
CHAT+
% control line ChAT+
ISL1+
% control line ISL1+
120
100
80
60
40
20
0
***
^^^
Con Con FALS SALS SALS
GAP19 - - - + - + - +
b
TUJ1+/CHAT-
% of control line Tuj1+/ChAT- cells
TUJ1+/ISL1-
% of control line Tuj1+/Isl1- cells
140
120
100
80
60
40
20
0
*
***
^^^
Con Con FALS SALS SALS
GAP19 - - - + - + - +

Fig.S8

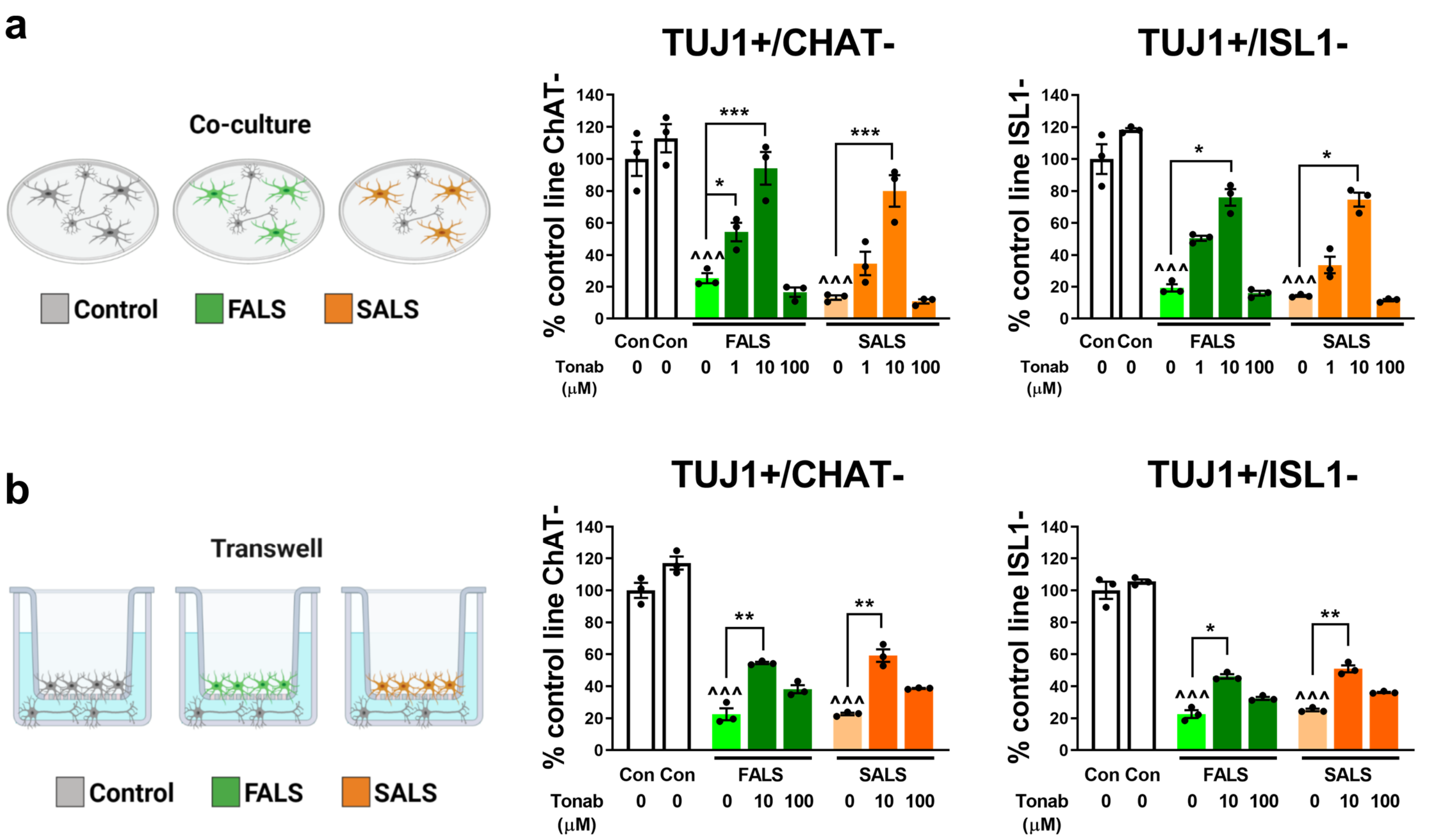
a
Co-culture
Control
FALS
SALS
TUJ1+/CHAT-
% control line ChAT-
TUJ1+/ISL1-
% control line ISL1-
Con Con
FALS
SALS
Tonab (μM)
0 0 0 1 10 100 0 1 10 100
b
Transwell
Control
FALS
SALS
TUJ1+/CHAT-
TUJ1+/ISL1-
Tonab (μM)
0 0 0 10 100 0 10 100

Fig.S9

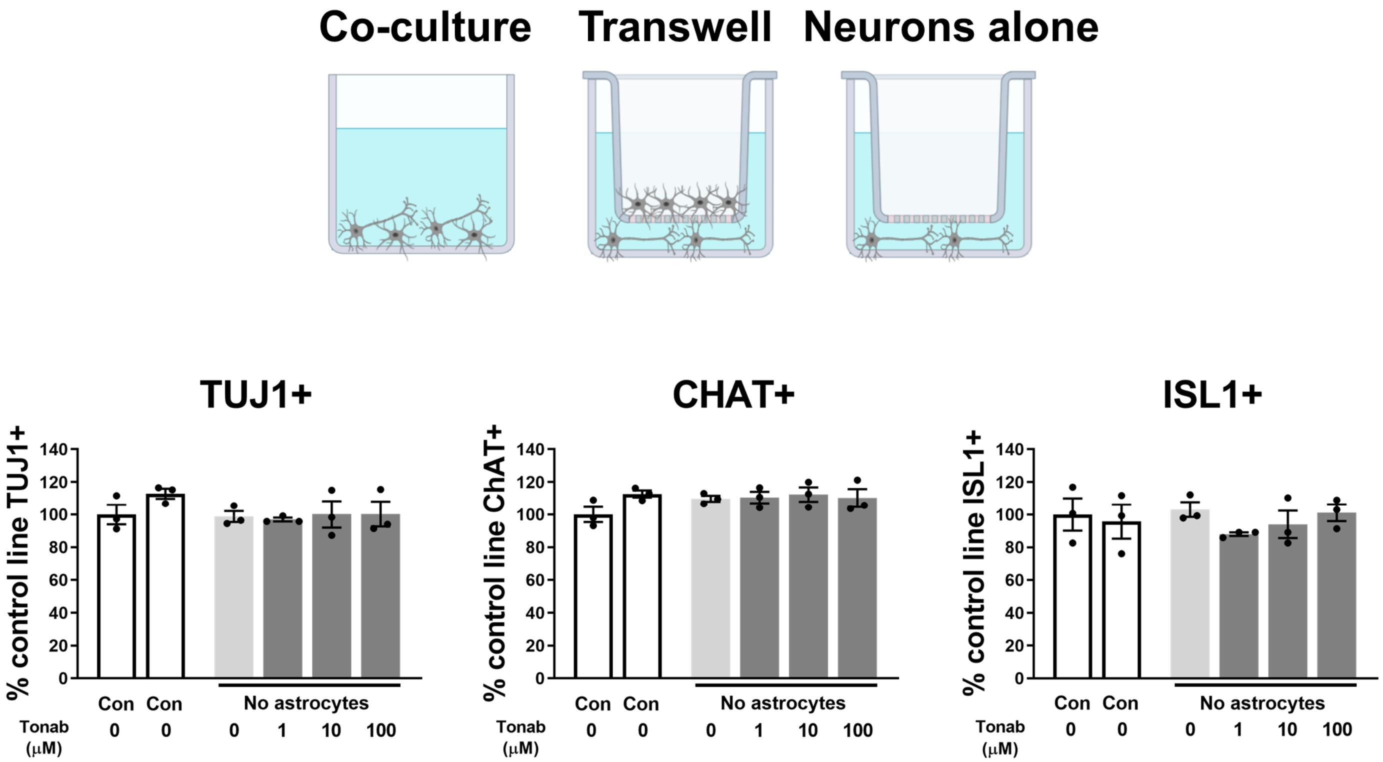
Co-culture
Transwell
Neurons alone
TUJ1+
% control line TUJ1+
CHAT+
% control line ChAT+
ISL1+
% control line ISL1+
Con
Con
No astrocytes
Tonab (μM)
0 0 0 1 10 100

Fig.S10

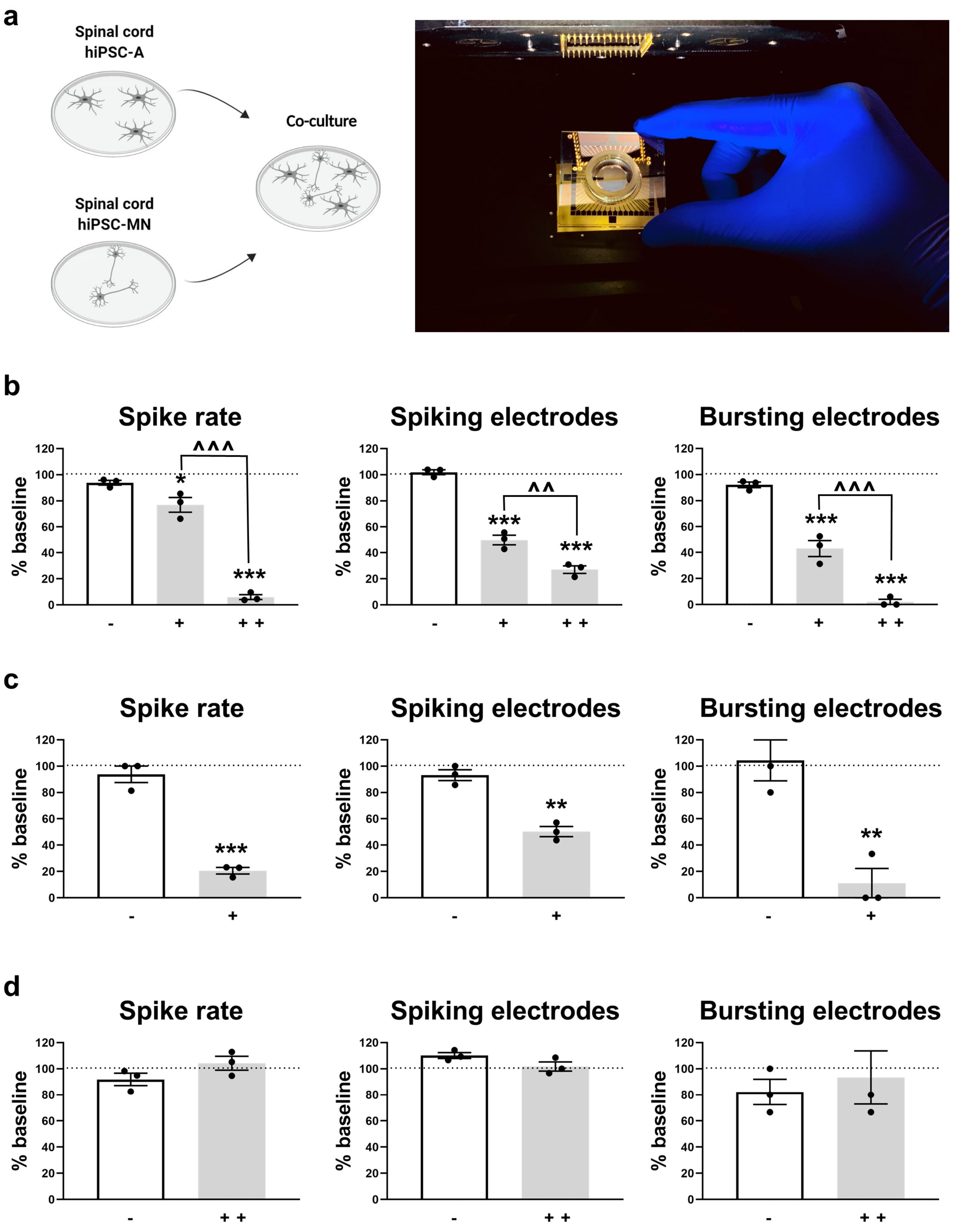
a
Spinal cord
hiPSC-A
Co-culture
Spinal cord
hiPSC-MN
b
Spike rate
Spiking electrodes
Bursting electrodes
% baseline
c
Spike rate
Spiking electrodes
Bursting electrodes
% baseline
d
Spike rate
Spiking electrodes
Bursting electrodes
% baseline

Fig.S11
